# Flexible and high-throughput simultaneous profiling of gene expression and chromatin accessibility in single cells

**DOI:** 10.1101/2024.02.26.581705

**Authors:** Volker Soltys, Moritz Peters, Dingwen Su, Marek Kučka, Yingguang Frank Chan

## Abstract

Gene regulation underpins development and is an intricate biological process involving transcription, typically at promoters within accessible chromatin. To understand cell-type specific regulatory networks, the ability to capture both transcription and chromatin accessibility simultaneously is crucial. However, joint measurements are technically challenging and current methodologies still face adoption challenges. Here, we present easySHARE-seq, an improvement on SHARE-seq, for the simultaneous measurement of ATAC- and RNA-seq in single cells. We address several limitations of the previous method by improving the barcode and streamlining the protocol. As a result, easySHARE-seq libraries have a usable sequence of up to 300bp (+200bp increase), making it suitable for e.g. investigation of allele-specific signals or variant discovery. Furthermore, easySHARE-seq libraries do not require a dedicated sequencing run thus saving costs. We applied easySHARE-seq to murine liver nuclei and recovered 19,664 nuclei with joint chromatin and expression profiles. By benchmarking against other combinatorial indexing-based techniques, we showed we can recover over 1.5 fold more transcripts per cell while retaining high scalability and low cost. To showcase our method, we identified cell types, exploited the multiomic measurements to link *cis*-regulatory elements to their target genes and investigated liver-specific micro-scale changes. We conclude that easySHARE-seq improves upon previous methods and can produce high-quality multiomic datasets. We expect it to be applicable to a wide range of study designs.

## Introduction

Development is an intricate biological process that requires the coordinated expression of thousands of genes, in millions of cells, nearly all of which carry the same genome, at least at the DNA basepair level. The difference between cells therefore emerges through regulation—acting before, during, and after transcription across layers of regulatory mechanism, resulting in lineage specification and proper tissue function^1,2^. Failures in these mechanisms underlie many human diseases: for instance, activating mutations in the Notch pathway can cause T-cell acute lymphoblastic leukaemia^3^. There are also many well-documented examples of disrupted enhancer–promoter communication contributing to congenital malformations and cancer^4–6^ as well as evolutionary adaptations^7,8^. Likewise, aberrant DNA methylation reshapes regulatory networks in malignancy^9^. For this reason, functional genomic methods that enable researchers to track developmental processes across regulatory layers are indispensable tools to help us understand gene regulatory processes in health and disease.

Over the past decade, large-scale initiatives such as ENCODE^10^ and FANTOM^11^ have expanded and spurred the development of a collection of molecular assays to interrogate many regulatory processes, including histone modifications^12^, open chromatin^13^, (nascent) transcription^14^, and three-dimensional genome conformation^15^ (for a detailed review, see ^16^). These approaches generally convert functional states into DNA sequence readouts and, together, have transformed our understanding of genome regulation^10^. These and other efforts to interrogate and summarize these results have converged on the idea where chromatin accessibility and long-range promoter–enhancer interactions are among the most predictive of gene expression programs and can bias cell fate decisions^2,17–19^.

Another inflection point in advances in sequencing and barcoding strategies is the increase in resolution of these functional assays, ultimately being able to distinguish individual cells from each other (“single-cell” resolution) ^20^. This change led to the discovery of rare cell populations and dynamic state transitions that were previously masked in conventional bulk assays. Consequently, a range of assays have been adopted to report on chromatin accessibility, transcriptomes, and protein abundance at the single-cell level—namely, scATAC-seq (for chromatin accessibility) ^21^, scRNA-seq (for the transcriptome) ^22^, and scBS-seq (for methylation) ^23^. Many of these approaches were based on physical separation between cells, classically in droplets. Paired with a correspondingly large set of DNA barcodes, this made it possible to pool thousands to now millions of cells together in a multiplexed sequencing reaction^24^. Yet, because each technology typically captures only one modality, inferring causal relationships between transcriptome and epigenome often relies on computational integration of separately measured datasets, using tools like Seurat^25^ or MOFA^26^. An inevitable assumption across these tools is that there is a shared underlying state across modality. While generally justifiable, it can also obscure fine-grained heterogeneity and complicate attempts at resolving how multiple regulatory layers interact within rare or transient cell types^27,28^.

Thus, the frontier is to measure multiple regulatory layers within the same single cell, enabling direct links between chromatin state and transcriptional output. Several strategies have been developed, each with distinct strengths and drawbacks. Early integrative methods such as scNMT-seq combined chromatin accessibility, DNA methylation, and transcription, but required technically demanding protocols and produced sparse coverage^27^. Generally, the early dependence on microfluidic chips and instrumentations imposed a limit on usability, throughput and protocol flexibility. These constraints encouraged the development of alternative approaches. One such approach that offers a simpler workflow and has found broad adoption is combinatorial indexing^29^. It relies on successive rounds of a “split-and-pool” procedure such that each cell is likely to be assigned a unique set of barcode segments. The sequential barcode design is powerful for single-cell barcoding as it scales exponentially and the “split-and-pool” procedure is more open to protocol improvements. Earlier examples include sci-CAR, which enabled joint profiling of chromatin accessibility and RNA, yet were limited by modest transcript capture and complex library design^30^. More recently, Paired-seq has been introduced as a paired multiomic approach, increasing capture efficiency for chromatin accessibility while maintaining scalability^31^. SHARE-seq improved throughput, robustness and data quality by refining barcoding strategies for simultaneous chromatin accessibility and transcriptional profiling. However, as with many protocols, its adoption has been constrained by custom sequencing requirements and extremely long barcode lengths^28^.

Commercial platforms have begun to address some of these barriers. The 10x Genomics Multiome assay offers a streamlined, kit-based solution for parallel scRNA-seq and scATAC-seq with standardized chemistry, but remains expensive and less adaptable to custom study designs. Scale Biosciences and Parse Biosciences provide a fully commercialized combinatorial indexing system that avoids microfluidics but currently does not offer simultaneous capture of scRNA- and scATAC-seq^32^. Collectively, these methods highlight the central technical challenges of joint profiling: first, balancing sensitivity and breadth across modalities; second, minimizing protocol complexity to ensure broad adoption; and last but not least, achieving scalability without compromising data quality. Addressing these limitations remains critical for charting how regulatory programs interact to specify cell fate and function.

Here, we describe an improved version of SHARE-seq with focus on minimizing protocol complexity and maximizing applicability. First, we redesigned the barcoding system, focusing on several improvements: 1. Reads can be sequenced up to a length of 300bp (instead of 100bp), allowing for variant discovery or assessment of allele-specific signals. 2. All libraries can be multiplexed with standard short-read libraries, decreasing sequencing costs. 3. Decreased protocol length by several hours. 4. The introduction of ‘sub-libraries’ allows for cost-effective sequencing of pilot data. We implemented these changes and applied them to a set of test samples to determine if easySHARE-seq can deliver larger and more flexible experimental designs through its more streamlined protocol and improved sequence quality.

## Results

### Technical improvements of easySHARE-seq

To develop easySHARE-seq, we implemented several improvements to SHARE-seq^28^ while retaining the broad overall workflow (**Fig. 1A**). First, we streamlined the barcode design. We retain a combinatorial indexing scheme but shorten each barcode segment to 7 nucleotides (nt), connected by 3nt overhangs (Hamming distance between barcode segments >= 2). This shortens the index sequences of the libraries to 17nt (from 99nt in SHARE-seq, Index1) and 8nt (Index2), respectively (**Fig. 1B**, **Suppl. Fig. 1A** shows a direct comparison). By using two segments of 192 barcodes each plus a separate third segment (to be added in a later PCR step), we achieve a total of 192×192×96 barcode combinations (~ 3.5 million) in the space of 25nt. The advantage of this design is that it allows more effective use of sequencingcycles and the sequencing of up to 300bp of the insert if desired (**Fig. 1C**; compared to 100nt in SHARE-seq or 90nt in Chromium Multiome libraries) while keeping the same scalability as SHARE-seq. Having longer insert read lengths increases the recovery of informative DNA variants (**Fig. 1C**), which is crucial in detecting allele-specific signals or cell-specific variant discovery, e.g., in F1 hybrids or cancer cells. Furthermore, this design allows simultaneous sequencing of easySHARE-seq libraries with standard Illumina libraries (‘multiplexing’), which can greatly reduce sequencing costs, especially in smaller pilot experiments. Separately, we shortened and optimized the bench protocol. The single-cell barcoding step is shortened from 4.5 h to 1.5 h by replacing all hybridization steps with a ligation round as well as eliminating blocking oligos. This has the advantage of minimizing library loss while improving yield. Additionally, after barcoding cells are aliquoted into as many ‘sub-libraries’ as desired, typically 3,500 cells each. This results in higher flexibility for the fine-tuning of cell numbers, the cost-effective sequencing of e.g. a pilot sub-library to assess overall data quality or the storing of sub-libraries for potential later use. All in all, easySHARE-seq reagents cost USD 0.056/cell, which is similar to the original SHARE-seq protocol. But, we anticipate much lower sequencing costs, as we now circumvent the highly customized 100nt index reads. For example, even a 50 cycle sequencing kit would result in sufficient information for many count-based applications. A comparison, including cost, between the two protocols as well as the most commonly used commercial solution can be found in **Table 1** (For a detailed description of the flexibility of easySHARE-seq, instructions on how to modify and incorporate the framework into new designs, how to convert it to a single-channel assay as well as critical steps to assess when planning to use easySHARE-seq see *Supplementary Notes*).

**Figure 1:**
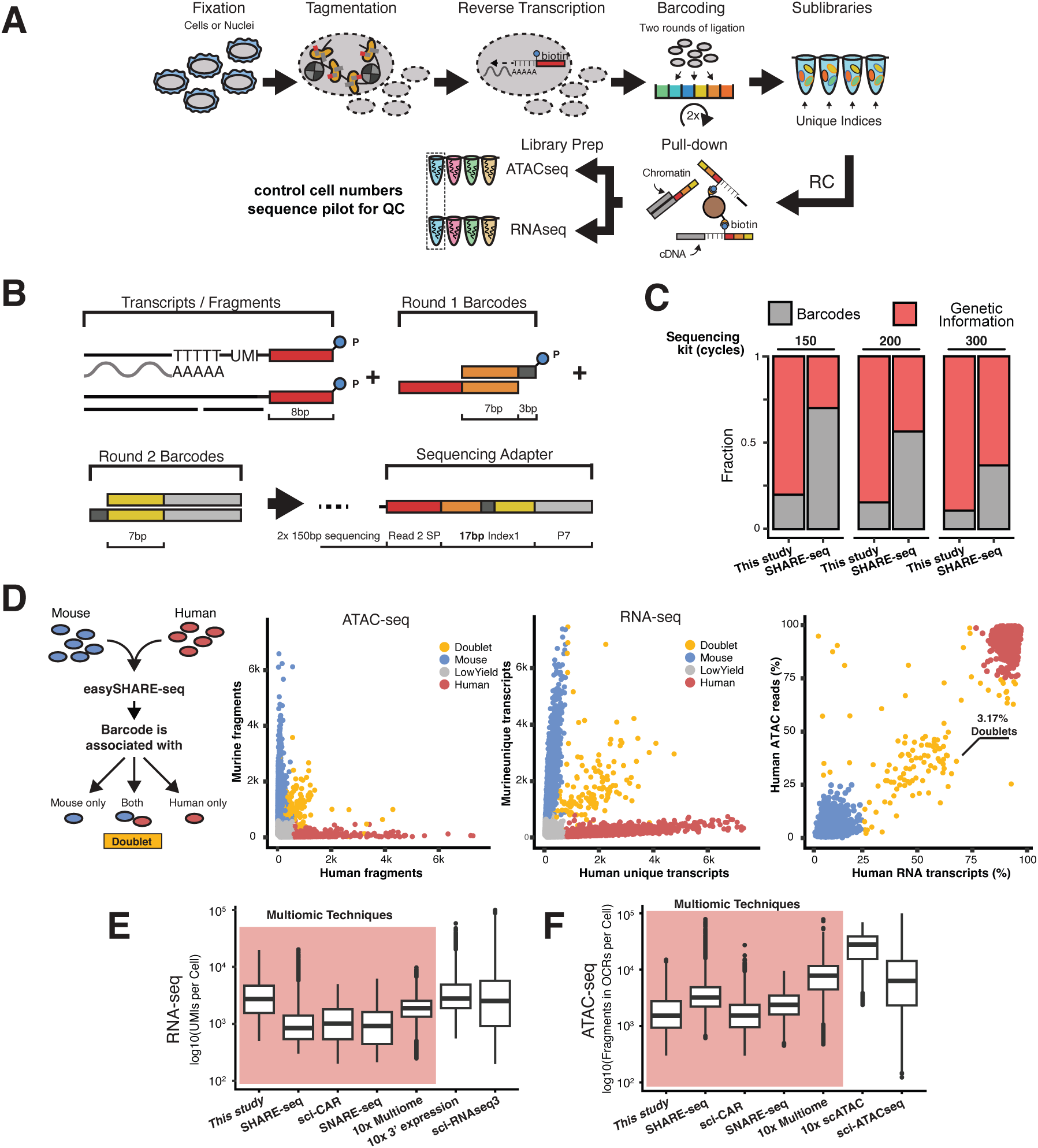
easySHARE-seq enables high-quality and accurate simultaneous scATAC-seq and scRNA-seq profiling. **(A)** Schematic workflow of easySHARE-seq. **(B)** Generation and structure of the cell-specific barcode within Index 1. Total length of the final barcode is just 17nt compared to 99nt previously. **(C)** Fraction of sequenced DNA bases allocated for either barcodes (grey) or genetic information (red) in easySHARE-seq and SHARE-seq using different sequencing kits. **(D)** Left: Principle of a species-mixing experiment. Murine OP-9 and human HEK cells are mixed prior to easySHARE-seq. After sequencing, sequences associated with each cell barcode are assessed for genome of origin. Middle left: Unique ATAC-seq fragments per cell aligning to the mouse or human genome. Cells are coloured according to their assigned origin (red: human; blue: mouse; orange: doublet). Middle right: Unique RNA-seq transcripts per cell aligning to the mouse or human genome. Right: Percentage of ATAC-seq fragments or RNA-seq transcripts per cell relative to total sequencing reads mapping uniquely to the human genome. 3.17% of all observed cells classified as doublets. Accounting for same-species doublets, this results in a doublet rate of 6.34%. **(E)** Comparison of UMIs/cell across different single-cell technologies. Red shading denotes all multiomic technologies. Datasets are this study, SHARE-seq ^28^ (murine skin cells), sci-CAR ^30^ (murine kidney nuclei), SNARE-seq ^33^ (adult & neonatal mouse cerebral cortex nuclei), 10x Multiome^34^ (murine liver nuclei), 10x 3’ Expression ^35^ (murine liver nuclei) and sci-RNAseq3 ^36^ (E16.5 mouse embryo nuclei). Cells have been downsampled to a common sequencing depth where possible (see Methods). **(F)** Comparison of fragments per cell across different single-cell technologies. Colouring as in (E). Datasets differing to (A) are 10x scATAC ^37^ (murine liver nuclei) and sciATAC-seq ^38^ (murine liver nuclei). Cells have been downsampled to a common sequencing depth where possible (see Methods).

**Table 1:** Comparison between multiomic single-cell technologies.

| Protocol | easySHARE-seq | SHARE-seq | 10x Multiome |
| --- | --- | --- | --- |
| Cost / Cell | 0.056\$ | 0.043\$ | 0.26\$ |
| Additional costs? | None | Dedicated sequencing run | Specialised Instrument |
| Throughput (cells) | > 200,000 | > 200,000 | 80,000 |
| Max. sequencing length of insert | 300bp | 100bp | 100bp |
| Processing time | ~12h | ~17h | ~14.5h |
| Convertible to single-channel assay? | Yes | Not described | No |

### easySHARE-seq labels both transcriptome and accessible chromatin in individual cells

To evaluate the accuracy and cell-specificity of the barcoding, we first performed easySHARE-seq on an equally mixed pool between human and murine cell lines (HEK and OP-9, respectively). Such a species-mixing design is a standard way to identify if a given single cell is consistently labelled with a single barcode^39^. Any barcode that comprises both human and murine reads is indicative of two or more cells sharing the same barcode (‘doublets’; **Fig. 1D**, left). In total, we recovered 3,808 cells (see *Methods* for details on cell filtering). After separately mapping the reads against the human and the mouse transcriptomes and genomes, we examined the proportions of each barcode’s reads that are mapped to the human or the mouse genomes.

Doing this, we found that both chromatin and transcriptome profiles separated well (**Fig. 1D**, middle). Overall, 96.83% of all cells (3,687) carried exclusively human or mouse chromatin and transcriptome. Between the modalities, we found that cDNA showed a lower accuracy with increasing transcript counts. This could be the result of lower mapping accuracy among transcripts, as coding regions are more conserved between species compared to intergenic regions^40^. Using the above criteria, we identified a total of 124 human–mouse doublets (3.17%; **Fig. 1D**, right). Factoring in undetected human–human and mouse–mouse doublets, we estimated a final doublet rate of 6.34%. For comparison, a 10X Chromium Next GEM experiment with 10,000 cells has a doublet rate of ~7.7% (reported by manufacturer). Importantly, easySHARE-seq doublet rates can easily be lowered further by aliquoting fewer cells within each sub-library, ensuring that there is sufficient barcode diversity to minimize barcode collision. We thus conclude that easySHARE-seq can support single-cell accurately measuring chromatin accessibility and gene expression in single cells.

### Performance of easySHARE-seq in murine primary liver cells

To assess data quality and overall performance of easySHARE-seq, we focused on murine liver. The liver consists of a diverse set of defined primary cell types, ranging from small non-parenchymal cell types such as Liver Sinusoidal Endothelial Cells^41^ (LSECs) to large hepatocytes, which potentially may be multinucleated. This provides a balance between tissue diversity and complexity for assessing our protocol. We generated matched high-quality joint chromatin and gene expression profiles for 19,664 adult liver nuclei across four age-matched mice (2 male, 2 female), for an estimated recovery rate of 70.2% (28,000 input cells). On average, we recovered 3,629 unique transcripts (± 2,990 transcripts, std) from 1,798 genes (± 969; **Suppl. Fig. 1E**). We used unique molecular identifiers (UMIs) to label individual transcripts, **Fig. 1B**; see *SI* for details). For chromatin accessibility, we recovered on average 2,213 unique fragments (± 1,995 fragments) from 2,011 accessible peaks (±1,672 peaks; **Suppl. Fig. 1E**). Closer examination of each data modality showed excellent recovery, which we will discuss in turn. For the transcriptome, 74% of total RNA-seq reads mapped to transcripts in a strand-specific manner (**Suppl. Fig. 1B**). We do note that 69.9% of reads mapped within introns (**Suppl. Fig. 1C**), which is consistent with the use of nuclei for transcriptome profiling. For comparison, this rate is lower than the ~75% intronic reads reported in Ma et al^28^. For chromatin accessibility, the scATAC-seq fraction of the libraries displayed the characteristic banding pattern during sample preparation (**Supp Fig. 1I**) and 55.9% of sequenced fragments are found in peaks on average (**Suppl. Fig. 1D**; ± 8.59%, range 30–76.2%). They are also highly enriched at transcription start sites (TSS; mean TSS Enrichment score 4.46; **Suppl. Fig. 1H**). For the relationship between sequencing depth and UMI/fragment recovery see **Suppl. Fig. 1F & G** and an example track can be seen at **Suppl. Fig. 1N**. For an estimate of ambient RNA contamination see **Suppl. Fig. 1O**. easySHARE-seq data was also highly reproducible, firstly across sub-libraries (R > 0.99) but also between biological replicates (R > 0.95; **Suppl. Fig. 1J & K**). These data gave us confidence that easySHARE-seq performed consistently across samples and batches and could form the basis of further in-depth analyses.

We next conducted benchmarking to determine how easySHARE-seq performed relative to other multiomic and representative single channel assays. To do so, we made use of publicly available datasets, where possible with matched sample type and controlling for sequencing depth by down-sampling. Most importantly, we used an identical analytical pipeline to allow proper comparison of results (**Fig. 1E & F**; note that down-sampling was not always possible due to lack of raw sequencing data, see Methods for a detailed description and figure legend for tissue type and study). We chose the following multiomic assays for benchmarking: SHARE-seq ^28^, sci-CAR ^30^, SNARE-seq ^33^ and 10x Multiome^34^. For added context, we also compared against single-modal datasets: for the transcriptome, 10X 3’ expression RNA-seq ^35^ and sci-RNAseq3 ^36^; and chromatin, 10X ATACseq ^37^ and sci-ATACseq ^38^. For the transcriptome, easySHARE-seq recovered significantly higher number of unique transcripts per cell (3,629 vs. 1,183, 1,302, 1,179 and 2,029 compared to SHARE-seq, sci-CAR, SNARE-seq and 10x Multiome Expression, respectively, *P* < 2×10^-16^ for all, pairwise t-test, two-tailed). As such, easySHARE-seq performed similarly to single modality assays albeit they still recovered significantly more transcripts per cell (3,901 and 4,607 for 10x and sciRNAseq3, respectively, *P* < 0.001). Summarized at the level of expressed genes, we also detected a higher number of genes expressed per cell (1,488 genes/cell vs. 608, 345, 708 and 1,222; *P* < 2×10^-16^; **Suppl. Fig. 1L**). Similarly, easySHARE-seq was closer to single-modal assays, even outperforming the 10X 3’ expression (1,343 for 10X and 2,010 for sciRNAseq3, *P* < 0.005). We hypothesize that the higher reported recovery from easySHARE-seq may have benefited from better sample retention as a result of protocol improvements, e.g., having fewer hybridization-and-wash steps, higher fixation, etc. These improvements could lead to increased UMI counts per gene (better signal dynamic range) and in turn, more genes passing threshold. However, since we were limited in the number of comparable publicly available dataset, we cannot rule out that the other benchmark datasets may not be fully comparable or representative. Also, despite our best efforts to make a fair comparison, our down-sampling procedure may not fully capture the complexity of the other datasets. Nevertheless, seeing that our dataset is ~2 times smaller than the other benchmarking datasets in terms of cells captured, we feel confident to conclude that easySHARE-seq is robust and flexible (via sub-libraries) and may offer an edge over other alternatives. Lastly, it is straightforward to adapt and upscale easySHARE-seq to a scRNA-seq protocol only (see *Supplementary Notes*).

Regarding ATAC-seq data quality, easySHARE-seq performed similarly to other published multiomic assays (**Fig. 1F**). but recovered less unique fragments per cell and detected less accessible peaks per cell than SHARE-seq (**Suppl. Fig. 1M**; median Fragments in peaks/cell: 1,329 (easySHARE-seq) & 3,220 (SHARE-seq)). This may be partially due to choosing to prioritise nuclei integrity and RNA-seq quality and therefore performing a higher fixation (0.35% PFA compared to 0.2% in SHARE-seq), which in our experience may result in less efficient transposition reactions. Nevertheless, 94.3% of open chromatin regions identified in this study overlapped those reported in the independent 10x Multiome dataset derived from the same tissue^34^ (**Fig. 1F**), providing cross-dataset validation of our ATAC-seq signal.

### Simultaneous scATAC-seq and scRNA-seq profiling in murine primary liver cells

To visualise and identify cell types, we projected the ATAC- and RNA-seq modalities separately into 2D Space and subsequently integrated modalities for a combined representation^42^ (**Fig. 2A**, **Suppl. Fig. 2 A&B**). Between the modalities, snRNA-seq showed good cluster separation, resulting in eight clusters (**Suppl. Fig. 2A**). Clustering in snATAC-seq alone resulted in only four major clusters, likely due to the lower dynamic range in chromatin accessibility and thus less information content (**Suppl. Fig. 2B**). We then annotated cell types on the integrated dataset based on gene expression of previously established marker genes^35,43^. Marker gene expression was highly specific to the clusters (**Fig. 2 B&C**, **Suppl. Fig. 2F**) and we recovered all expected cell types (**Suppl. Fig. 2C**, 83% Hepatocytes, 7.9% LSECs, 3% Hepatic Stellate Cells (HSCs), 2.04% Kupffer Cells, 2.01% B Cells, 1.2% Neurons, 0.4% Monocytes and 0.3% Cholangiocytes). Data recovery per cell differed between cell types, with LSECs having the least UMIs and fragments per cell on average (mean 2,644 UMIs/cell and mean 1,589 fragments/cell) whereas HSCs had the highest amount of transcripts (mean 3,764 UMIs/cell) and Neurons the most fragments (mean 3,040 fragments/cell; **Suppl. Fig. 2D & E**).

**Figure 2:**
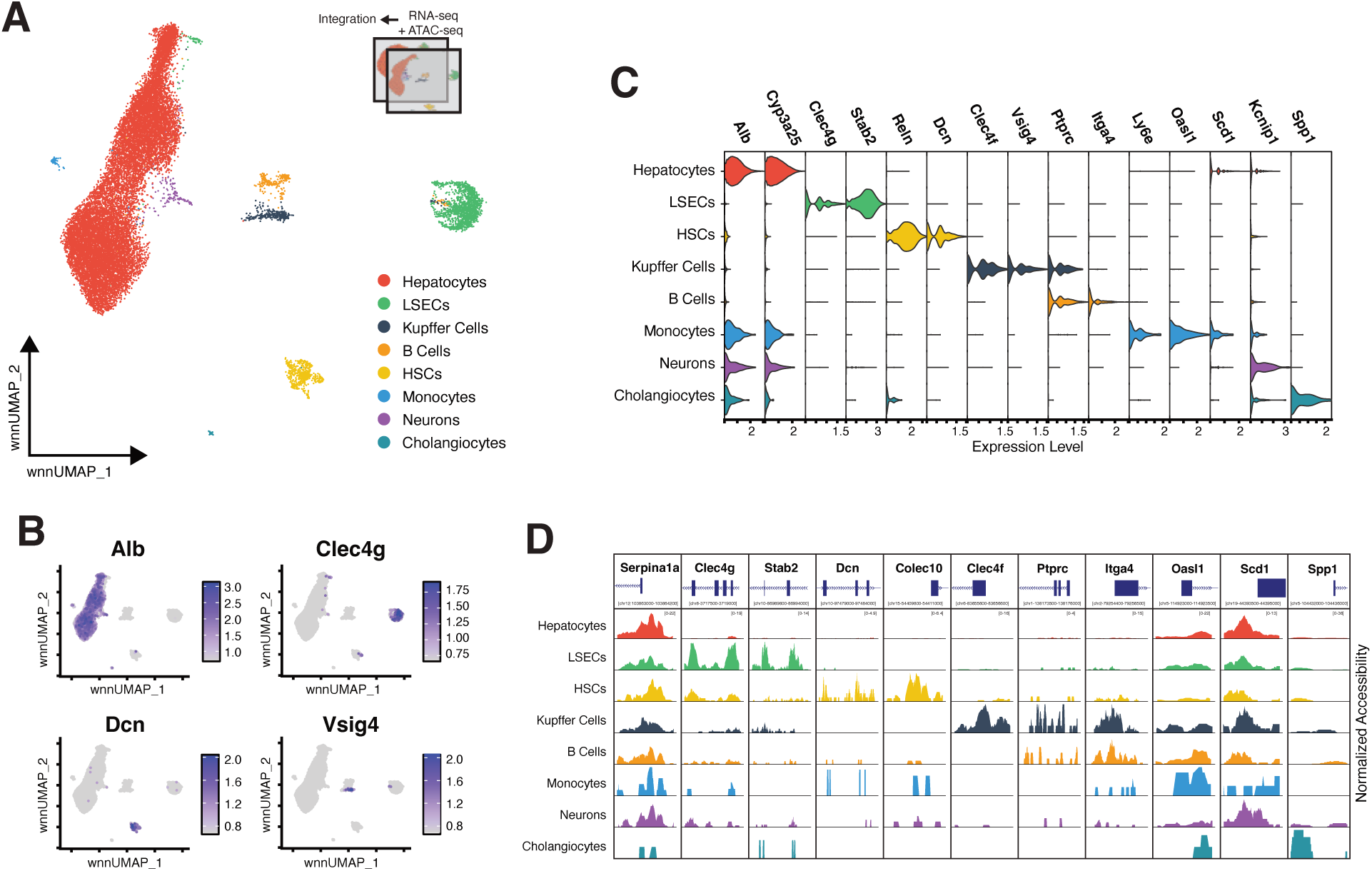
Cell type classification in primary liver nuclei using joint expression and chromatin accessibility profiles. **(A)** UMAP visualisation of WNN-integrated scRNA-seq and scATAC-seq modalities of 19,664 liver nuclei. Nuclei are coloured by their assigned cell type identity. **(B)** WNN-UMAPs of 19,664 liver nuclei with nuclei coloured according to the mean expression strength of marker genes for several cell types (*Alb*: Hepatocytes, *Clec4g*: Liver Sinusoidal Endothelial Cells (LSECs), *Dcn*: Hepatic Stellate Cells (HSCs), *Vsig4*: Kupffer Cells). Plots for further marker genes can be found in **Suppl. Fig. 2F**. **(C)** Violin Plots depicting the distribution of the normalised expression level of marker genes in all cells assigned to each cell type. **(D)** Pseudo-bulk ATAC-seq coverage tracks of normalised chromatin accessibility in all cells of each cell type in open chromatin regions overlapping marker genes.

Examining our dataset, we also found diversity within cell types. In the liver, hepatocytes fulfill different metabolic functions depending on their physical location in their organising structure called lobules (‘zonation’). These differences in function result in gene expression gradients along zonation, which are well described in hepatocytes^34,44^. This diversity was also reflected in our dataset (**Suppl. Fig. 2F**). For example, zonation markers such as *Gls2*, *Glul* and *Cyp3e1* expression showed clear gradients within hepatocytes.

Lastly, examining pseudo-bulk ATAC-seq tracks of the identified cell types showed highly cell-type specific signals despite lower clustering resolution, indicating that cell-type specific signals can be identified using each modality independently and showcasing the high congruence between the scATAC-seq and scRNA-seq modalities (**Fig. 2D**). Altogether, our results show that easySHARE-seq generates high quality and reproducible joint cellular profiles of chromatin accessibility and gene expression within primary tissue, expanding our toolkit of multiomic protocols.

### Uncovering the cis-regulatory landscape of key regulators through peak-gene associations

The power of multiomic technologies lies in simultaneously capturing information from multiple molecular layers, providing a more holistic view of the underlying biology. For example, connecting a cis-regulatory element (CREs) to its target gene is difficult with separate measurements even though it represents the pivotal step in transcription initiation^1^. However, changes to such relationships can impact gene function, and in many cases, cause diseases^45–47^.

To demonstrate the ability of easySHARE-seq data to support such analyses, we focused on LSECs (1,561 total), the cell type with the lowest amount of transcripts and fragments in our dataset (**Suppl. Fig. 2D & E**). Following Ma et al.^28^, we computed the correlation between gene expression and chromatin accessibility at nearby peaks to identify these putative CREs (pCREs, **Fig. 3A**, testing all accessible peaks within ± 500kb of the TSS and controlling for GC content and accessibility strength). In total, we found 81,514 significant peak–gene associations (45% of total peaks, *P* < 0.05) with 15,061 genes having at least one association (65.8% of all genes; **Suppl. Fig. 3A & C**). In rare cases (2.9%), these pCREs were associated with five or more genes. This drops to 0.03% when considering only pCREs within ± 50 kb of a TSS (**Suppl. Fig. 3C & D**). These pCREs tended to cluster to regions of higher expressed gene density (2.15 mean expressed genes within 50kbp vs 0.93 for all global peaks) and their associated genes were enriched for biological processes such as mRNA processing, histone modifications and splicing (**Suppl. Fig. 3H**), possibly reflecting loci with increased regulatory activity. Although potentially informative, these signals may reflect technical artifacts. We therefore restricted the analysis to the most significant gene association per peak, resulting in 40,975 pCREs. Exploring these associations, we found that linked genes exhibit significantly higher expression levels and linked peaks greater chromatin accessibility relative to their non-linked counterparts (**Suppl. Fig. 3 I,J**) which could either reflect increased power to detect them or suggest that cis-regulatory associations are enriched at transcriptionally more active loci. Despite this observation, chromatin accessibility and gene expression were uncorrelated within linked pairs (**Suppl. Fig. 3K**; p = 0.39, r=0.004). This indicates that chromatin accessibility poorly predicts enhancer activity, consistent with evidence that promoters integrate enhancer signals in a non-linear fashion^48,49^.

**Figure 3:**
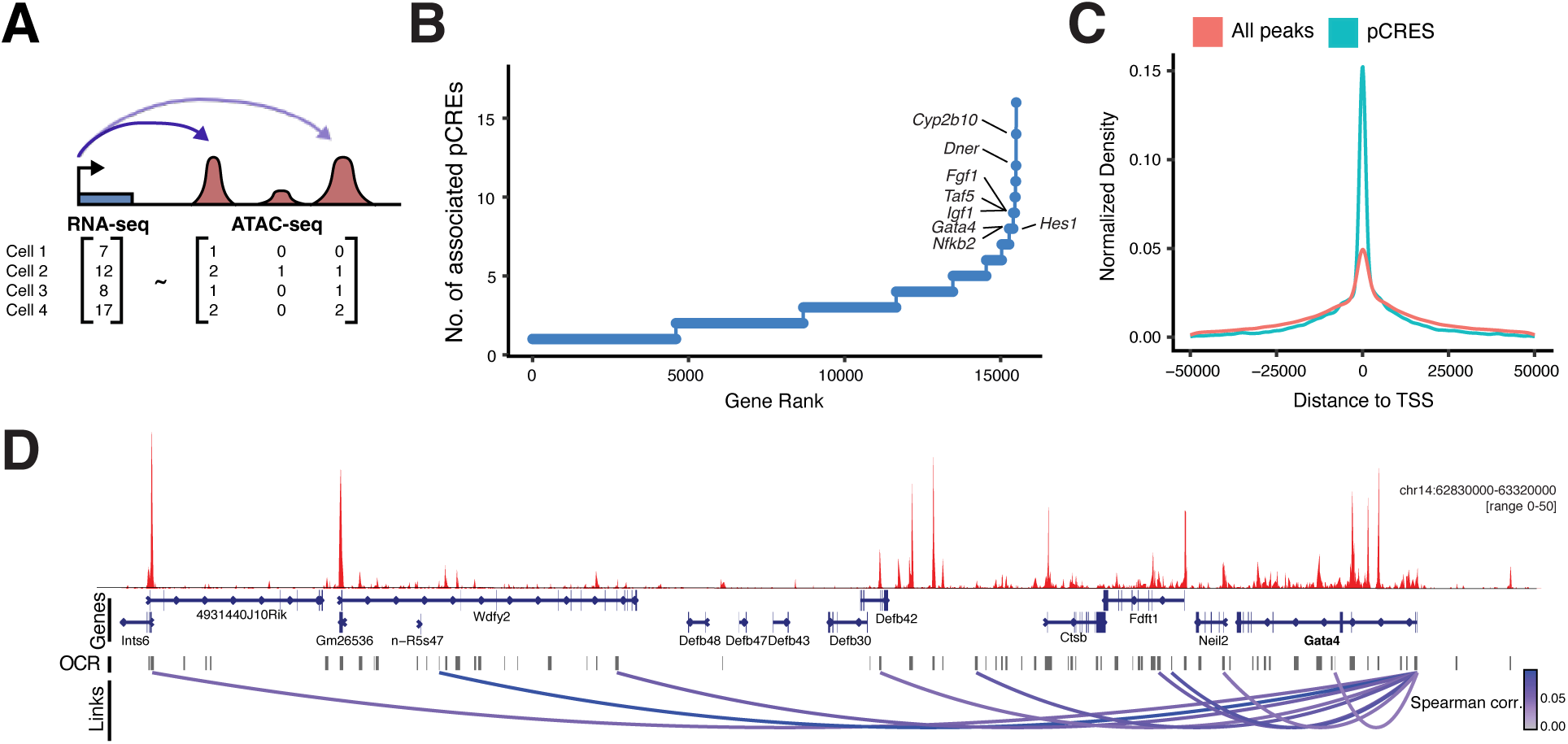
Assigning putative cis-regulatory elements to their target genes by correlating simultaneous measurements of gene expression and chromatin accessibility. **(A)** Schematic depicting the concept for linking putative cis-regulatory elements (pCREs) to their target genes. For each gene, all open chromatin regions (OCRs) within +− 500bk of its transcription start site (TSS) are tested. A pCRE is linked to a gene if the spearman-correlation of its chromatin accessibility to the gene’s expression falls outside the expected distribution estimated by correlating chromatin accessibility of 100 unrelated peaks (on different chromosomes) to the gene expression. (**B**) Genes ranked by their number of significantly correlated pCREs (*P* < 0.05, ±500kbp from TSS) in Liver Sinusoidal Endothelial Cells (LSECs). Marked are genes in the top 1% that are either transcription factors or cell-type specific regulators shown to fulfill a critical role in LSECs. (**C**) pCREs are enriched for TSS proximity. Normalised density of all open chromatin regions within ±50kbp of a TSS (red) and of all pCREs within ±50kbp of a TSS (blue). (**D**) Aggregate snATAC-seq pileup (red) of LSECs at the *Gata4* locus and 500kbp upstream region. Grey bars indicate open chromatin regions. Loops denote pCREs significantly correlated with *Gata4* and are coloured by Spearman correlation of respective pCRE–Gata4 comparison.

We then ranked genes based on their number of associated pCREs (**Fig. 3B**). Within the top 1% of genes with the most pCRE associations were many key regulators and transcription factors. Examples include *Taf5*, which directly binds the TATA-box^50^ and is required for initiation of transcription, or *Gata4*, which has been identified as the master regulator for LSEC specification during development. For instance, LSEC-specific knock-out of *Gata4* in adult or embryonic mice both lead to transdifferentiation of discontinuous LSEC into continuous capillaries, resulting in liver hypoplasia and fibrosis, even lethality in the case of embryonic conditional loss-of-function^51,52^. In adult mice, LSECs also control regeneration and metabolic maturation of liver tissue in adult mice. As such, it incorporates a variety of signals and its expression needs to be strictly regulated, which can be related to the many pCREs associations (8 total; **Fig. 3D**). Similarly, *Igf1* also integrates signals from many different pCREs^53^ (9 total; **Suppl. Fig. 3G**). Notably, pCREs are significantly enriched at transcription start sites (TSS), even relative to background enrichment in global peaks (**Fig. 3C**). We thus conclude that easySHARE-seq makes it possible to directly investigate the relationship between chromatin accessibility and gene expression and link putative cis-regulatory elements to their target genes at genomic scale, even in relatively rare cell types with low mRNA contents.

### De Novo identification of open chromatin regions and genes displaying zonation in LSECs

Micro-scale changes in gene regulation or expression can often have a major impact, not only within a cell through cell-fate determination but also via cell–cell communications, for example during embryonic development or in certain diseases^7,54^. To demonstrate the ability of easySHARE-seq to capture and assess these micro-scale changes, we investigated the process of zonation in LSECs. The liver consists of hexagonal units called lobules where blood flows from the portal vein and arteries toward a central vein^55,56^ (**Fig. 4A**). The central–portal (CP) axis is characterised by a morphogen gradient, e.g. *Wnt2*, secreted by central vein LSECs and thus activating the canonical *Wnt* pathway, giving rise to spatial division of labour among cells along it (‘zonation’)^57–59^. While studying zonation in hepatocytes is more common, doing so in non-parenchymal cells such as LSECs is challenging as these are physically small cells with low mRNA content (**Suppl. Fig. 2D & E**), often lying below the detection limit of current spatial transcriptomic techniques. As a result, only very few studies assess zonation in LSECs on a genomic level^60^. However, LSECs are critical to liver function as they line the artery walls, clear and process endotoxins. As outlined above in discussing its master regulator *Gata4*, LSECs are necessary for proper liver regeneration and setting up morphogen gradients to regulate hepatocyte gene expression^61–63^. For these reasons, a fine-grained understanding of these cells and their gene regulation is crucial for tackling many diseases.

**Figure 4:**
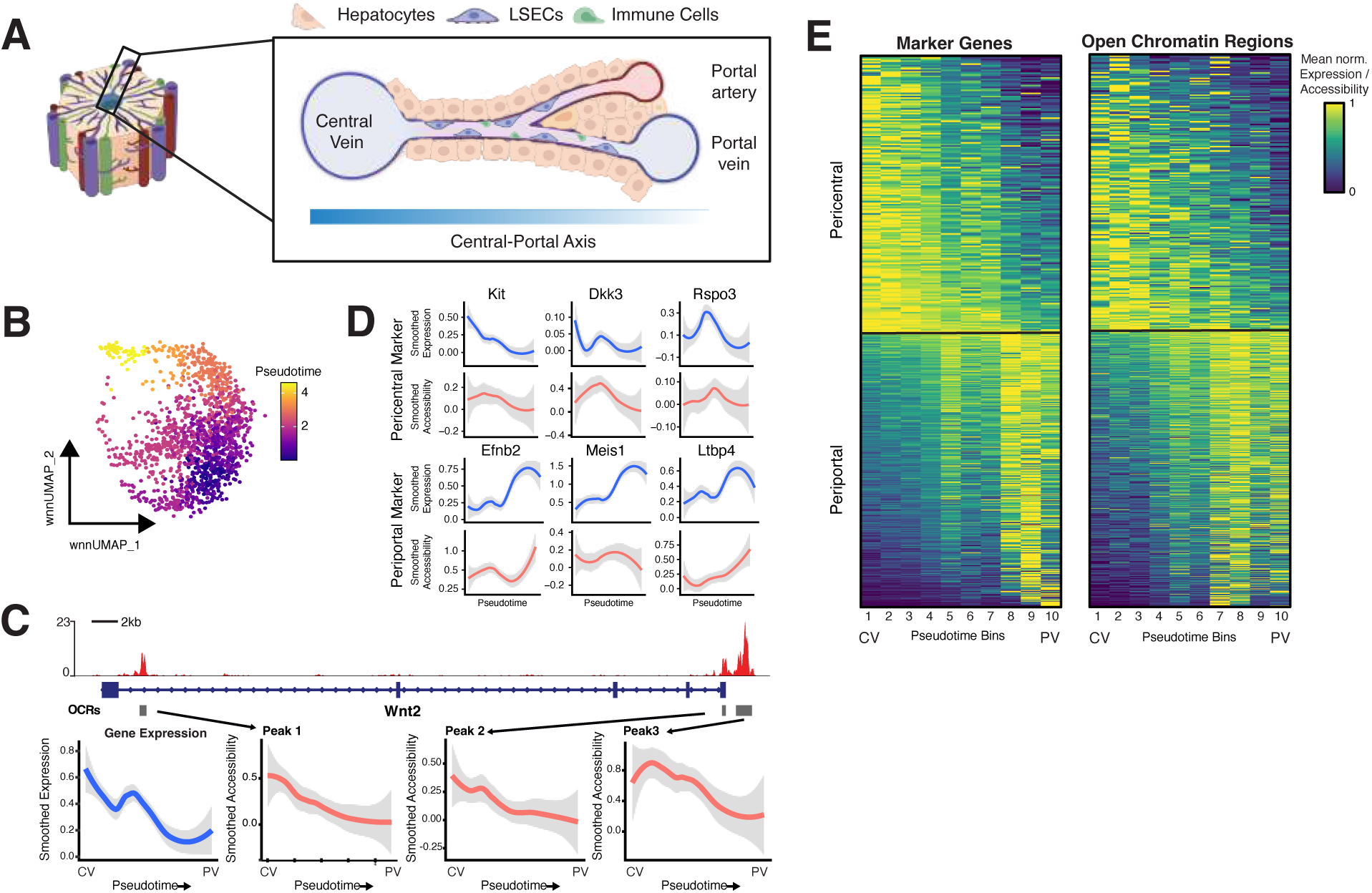
Zonation profiles in LSECs across gene expression and chromatin accessibility. **(A)** Schematic depiction of a liver lobule. A liver lobule has a ‘Central–Portal Axis’ defined by morphogen gradients starting from the central vein to the portal vein and portal artery. The sinusoidal capillary channels are lined with Liver Sinusoidal Endothelial Cells (LSECs). (**B**) UMAP of 1,561 LSECs coloured by pseudotime. (**C**) Changes along the Central–Portal Axis at the *Wnt2* locus. Top: Aggregate snATAC-seq profile (red) of LSECs at the *Wnt2* locus. Grey bars denote identified open chromatin regions (OCRs). Bottom: In blue, loess trend line of mean normalised *Wnt2* gene expression along pseudotime / the Central–Portal-Axis (CP-axis, central vein, CV; portal vein, PV). In red, loess trend line of mean normalised chromatin accessibility in OCRs at the *Wnt2* locus along the CP-axis. (**D**) Loess trend line of mean normalised gene expression (blue) of marker genes and mean normalised chromatin accessibility (red) at OCRs overlapping the marker gene along the CP-axis for pericentral markers (top, increased toward the central vein, *Kit*, *Dkk3* and *Rspo3*) and periportal markers (increased toward the portal vein, *Efnb2*, *Meis1* & *Ltbp4*) (**E**) Left: Zonation profiles of 550 genes along the CP axis. Right: Zonation profiles of 744 open chromatin regions along the CP axis. All profiles are normalised by their maximum.

We therefore asked if we can recover zonation gradients and potentially identify novel marker genes and open chromatin regions displaying zonation from our data. Specifically, we ordered LSECs along pseudotime and then checked marker gene expression and chromatin accessibility gradients along it, asking if this recapitulates zonation gradients discovered using Paired-cell sequencing (**Fig. 4B-D**)^60^. This analysis recovered the expected zonation profiles for several known zonation markers. For example, *Wnt2* expression decreased strongly along the CP axis as did chromatin accessibility of all three peaks at the *Wnt2* locus (**Fig. 4C**). We also recovered the expected zonation profiles for several further known pericentral (decrease along the CP-axis, e.g. *Rspo3*, another canonical *Wnt* agonist & *Kit*) and periportal (increase along the CP-axis, e.g. *Efnb2* & *Meis1*) marker genes as well as chromatin accessibility gradients at their associated open chromatin regions (**Fig. 4D**). Knowing that the imputed pseudotime faithfully ordered cells along their zonation gradients, we sought to identify novel genes and open chromatin regions displaying zonation in LSECs based on the decrease or increase of the rolling mean in expression or accessibility along pseudotime (see Methods). In total, we classified 179 genes and 198 open chromatin regions as pericentral and 371 genes and 546 open chromatin regions showed periportal zonation profiles (**Fig. 4E**). The list of markers contained many genes regulating epithelial growth and angiogenesis (e.g. *Efna1, Nrg2, Jag1, Bmp2, Bmper, Nrp2*)^64–67^, related to regulating hepatocyte functions and communication (e.g. *Dpp4, Foxo1, Foxp1, Insr, Ephb4*)^68–71^ as well as immunological functions (e.g. *Il1a, Pigr, Tgfb1*)^72,73^, suggesting that these processes show variation along the PC axis. As many of these newly identified markers are implicated in liver illnesses such as cirrhosis, fibrosis or non-alcoholic fatty liver disease^74,75^, these genes are potential new biomarkers for their identification and the open chromatin regions starting points for investigating the role of gene regulation in their emergence. This demonstrates the sensitivity and applicability of easySHARE-seq to detect micro-scale changes even in lowly expressed cell types and simultaneously underscores the advantage of multiomic measurements, where pseudotime inferred from gene expression can then be linked to chromatin state dynamics.

## Discussion

Understanding complex processes such as gene regulation or disease states requires the integration of multiple layers of information. To this end we have developed easySHARE-seq to streamline the generation of high-quality joint profiling of chromatin accessibility and gene expression within single cells in a way that can accommodate diverse experimental designs. In this section, we will discuss the following topics in turn: the improvements, the implications on experimental designs, current limitations and further improvements.

### Protocol improvements

We noted that despite the substantial impact from the original SHARE-seq publication, there has not been widespread adoption of the protocol itself by other groups. One possible explanation could be the sheer number of customizations required (ordering and building the original barcodes, custom sequencing runs, etc.), which poses a high barrier of entry for many other users. Taking these into account, we have incorporated a number of improvements in developing easySHARE-seq: barcode simplification, reallocation of read lengths, protocol streamlining and the ability to split a larger sample into “sub-libraries”.

The most consequential change in the protocol was our redesign of the barcodes. Here, we have optimized for barcode complexity within a short sequence space, while preserving sufficient molecular stability throughout the multiple rounds of handling and buffer changes. The species-mixing experiment, together with the high consistency in cell-type specific signals gave us confidence that our strategy was sufficiently robust for multi-modal single-cell profiling. Having significantly shortened the barcodes (from 107nt to 25nt), we could now retain much of the read lengths for the transcripts or fragments themselves (in other words: data). We argue that just read counts in transcriptome or chromatin accessibility profiling capture only some—but far from all—aspects of gene regulation. There are entire aspects of regulatory mechanisms that benefit from longer read lengths such as variant detection and subsequent analysis of allele-specific signals, RNA-editing, insertion/deletions in CREs and more. These applications may be entirely missed under the original SHARE-seq configuration. Next, a number of our protocol improvements, when taken in combination, go beyond their individual effects. For instance, by shortening the protocol (12 vs. 17 h) and streamlining various enzymatic steps, we showed that we can achieve better sensitivity in RNA-seq (greater UMIs per cell, among others; Fig. **1E**). This, together with our introduction of sub-libraries using the i5 barcoding segment, enabled both higher throughput and greater flexibility in experimental design. This in turn allows users to conduct smaller-scale pilot tests, while retaining the option to increase sequencing efforts after confirming the quality of the libraries. In our experience, such flexibility is crucial in day-to-day experimental designs and is often not captured or not possible in previous protocols. In terms of costs per cell, easySHARE-seq performs similarly to standard SHARE-seq with ~5.6 cents/cell, which is before factoring in the reduction in sequencing costs compared to standard SHARE-seq, and a fraction of the costs (<25%) of commercially available platforms without considering their specialised instrument costs.

### Experimental designs

As alluded to in the last section, we think easySHARE-seq holds a number of technical advantages that translate well into an ability to support more (and different) experimental designs. For example, having the longer read-lengths and thus better power to resolve genomic variants will help in the majority of cases where the samples are diverse, from non-inbred individuals or may carry de novo mutations as in cancer. Having sub-libraries in combination with shorter experimental time makes easySHARE-seq more practical when performing multiple experiments, e.g. assaying more conditions or biological replicates, as in practice, one big experiment with hundreds of thousands of cells rarely reflects actual experimental design.

Our results from the murine LSECs also show that we can recover cells that may be rare or have very low mRNA content. This in turn implies that we may be able to use easySHARE-seq when investigating diseases originating from rare cell populations. For example, pancreatic ß-cells make up only 1-2% of the endocrine tissue in the pancreas yet their failure to function has major implications for human health^76^. As demonstrated in the example of LSEC zonation and *Gata4*, the key cell–cell signaling hub during cell specification may involve specific cell populations to set up morphogen gradients like *Wnt2*, or specific genes like *Gata4* to initiate regulatory cascades. In cases like this, researchers may benefit from having access to both regulatory channels, as solely focusing on gene expression or chromatin accessibility alone may not reveal the true significance of the shift. Another area where this is potentially beneficial is in evolutionary studies. For example, this allows exploring how gene-enhancer dynamics change between species or populations and how that in turn might shape differences in the transcriptome.

### Limitations and future developments

While we are encouraged by the improvements we have implemented in easySHARE-seq, there are still a number of limitations that will require future work to address. For example, single-channel assays still produce higher quality data compared to our multiomic protocol. Additionally, validating this protocol in a widely used cell line would provide independent confirmation of its improvements. Second, there is opportunity to increase ATAC-seq data quality further without compromising the RNA-seq channel as compared to SHARE-seq, it somewhat lacks resolution. Potential experimental steps that can be optimized are different aspects of fixation or tagmentation; for a detailed description see *Supplementary Notes*. We acknowledge that the current relatively lower quality of the ATAC-seq data may introduce increased variance in downstream analyses, particularly in the context of multi-modal integration with gene expression data making data interpretation more challenging. Lastly, while easySHARE-seq is significantly less expensive than 10x or SHARE-seq (factoring in sequencing costs), it still requires a significant initial investment for the DNA oligos.

In terms of future developments of this assay, other potential improvements might be the introduction of sample-specific barcodes to allow multiplexing many samples at once. While in the RNA-seq, this can be achieved rather easily (via the RT-primer, see *Supplementary Notes*), this is more challenging in the ATAC-seq. Furthermore, there is potential to decrease the amount of reagents such as T4 Ligase thus decreasing overall costs and increasing barcode complexity further to handle millions of cells simultaneously.

## Conclusion

In conclusion, we present here easySHARE-seq, including benchmarking against existing methods and real-life examples. While the presented biological examples may be more limited in scope, it contains a number of cell types, including examples of micro-differentiation that goes some way towards demonstrating that easySHARE-seq is not only applicable to a variety of study designs but also that the simultaneous measurements can be used to disentangle the complex molecular hierarchy and relationships between different regulatory layers. We envision easySHARE-seq as an important technological step toward the more widespread and diverse use of multiomic techniques. Ultimately, these techniques will play an important role in understanding gene regulation in health and disease, differentiation or lineage commitment, and determining genetic variants affecting those processes.

## Supporting information

LSECs Peak-Gene Associations

LSECs Gene Zonation Markers

LSECs Peaks Zonation Markers

Celltype Annotations

---

**Supplementary Figure 1:**
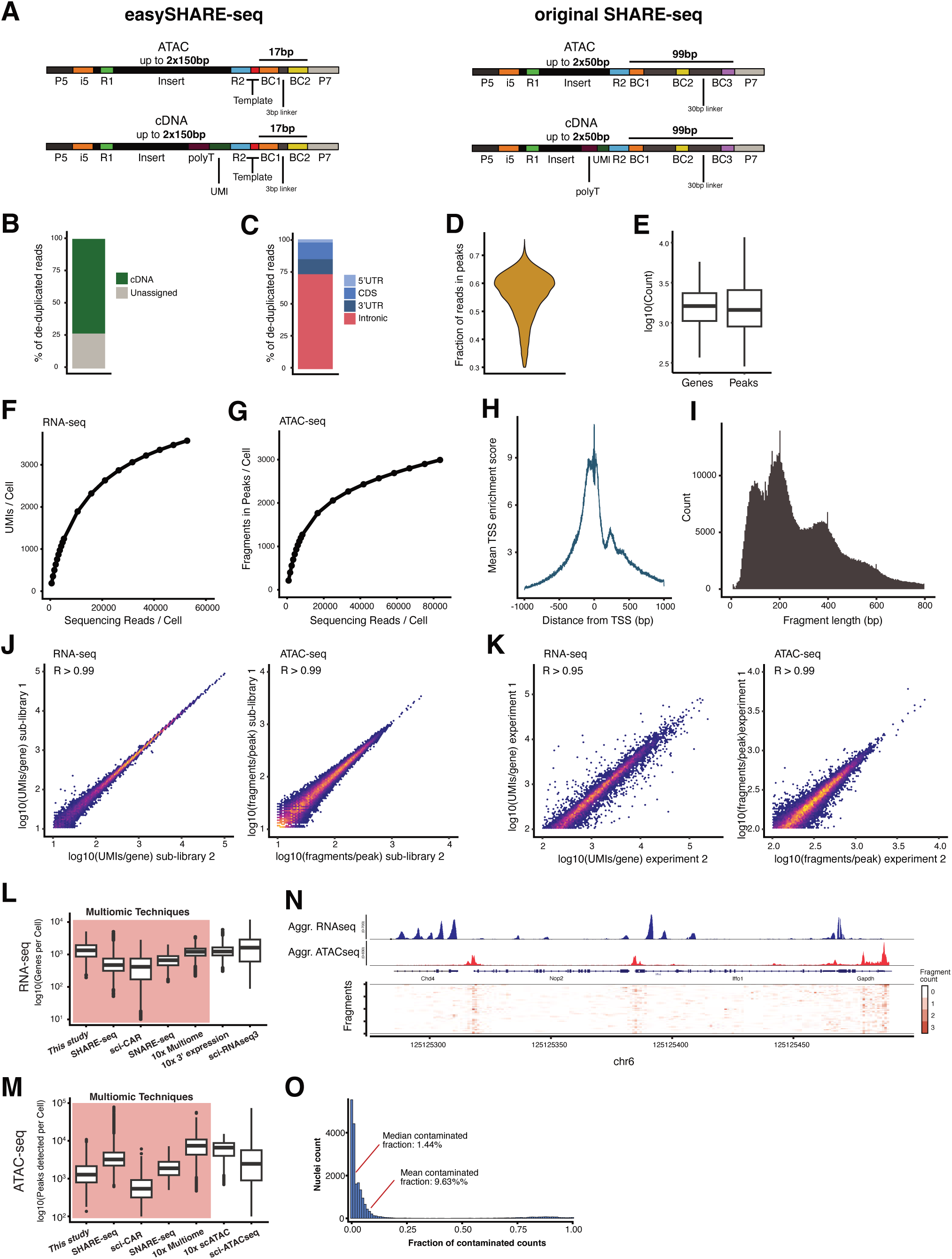
Barcode structure and summary of quality control measures in liver nuclei. **(A)** Structure of scATAC-seq and scRNA-seq sequencing reads in easySHARE-seq and the original protocol (BC1/2: barcode segment 1/2, UMI: Unique Molecular Identifier. (**B**) Percentage of total scRNA-seq sequencing reads containing cDNA fragments in murine primary liver nuclei. (**C**)Percentage of de-duplicated scRNA-seq sequencing reads overlapping an exon, intron, 5’UTR or 3’UTR. (**D**) Distribution of fraction of reads in peaks (FRiP) per cell in scATAC-seq data in murine primary liver nuclei (mean FRiP: 0.55). (**E**) Boxplot depicting the distribution of expressed genes and accessible peaks per cell in murine primary liver nuclei (mean expressed genes: 1,798; mean accessible peaks: 1,983) (**F**) Number of mean UMIs per cell recovered in the snRNA-seq when subsampling to different raw sequencing depths. (**G**) Number of mean fragments in peaks per cell recovered in the snATAC-seq when subsampling to different raw sequencing depths. (**H**) Mean transcription start site (TSS) enrichment score per cell in relation to distance from nearest TSS in the snATAC-seq data. (**I**) Histogram of fragment length in snATAC-seq sequencing reads. (**J**) Reproducibility of easySHARE-seq between sub-libraries shown by comparing the number of UMIs recovered per gene or peak across them. Each dot depicts either a gene (left) or peak (right). (**K**) Reproducibility of easySHARE-seq between biological replicates shown by comparing the number of UMIs recovered per gene or peak across biological replicates. Each dot depicts either a gene (left) or peak (right). (**L**) Comparison of genes expressed per cell across different single-cell technologies. Red shading denotes all multiomic technologies. Datasets are the same as in Fig. 1E. Cells have been downsampled to a common sequencing depth where possible (see Methods). (**M**) Comparison of accessible peaks per cell across different single-cell technologies. Colouring as in (**L**). Datasets are the same as in Fig. 1F. Cells have been downsampled to a common sequencing depth where possible (see *Methods*). (**N**) Aggregated scRNA- and scATAC-seq tracks of easySHARE-seq at the GAPDH locus. (**O**) Histogram of fraction of contaminated RNA counts (UMIs) per nuclei as estimated by decontX. For a discussion of ambient RNA contamination see *Supplementary Notes*.

**Supplementary Figure 2:**
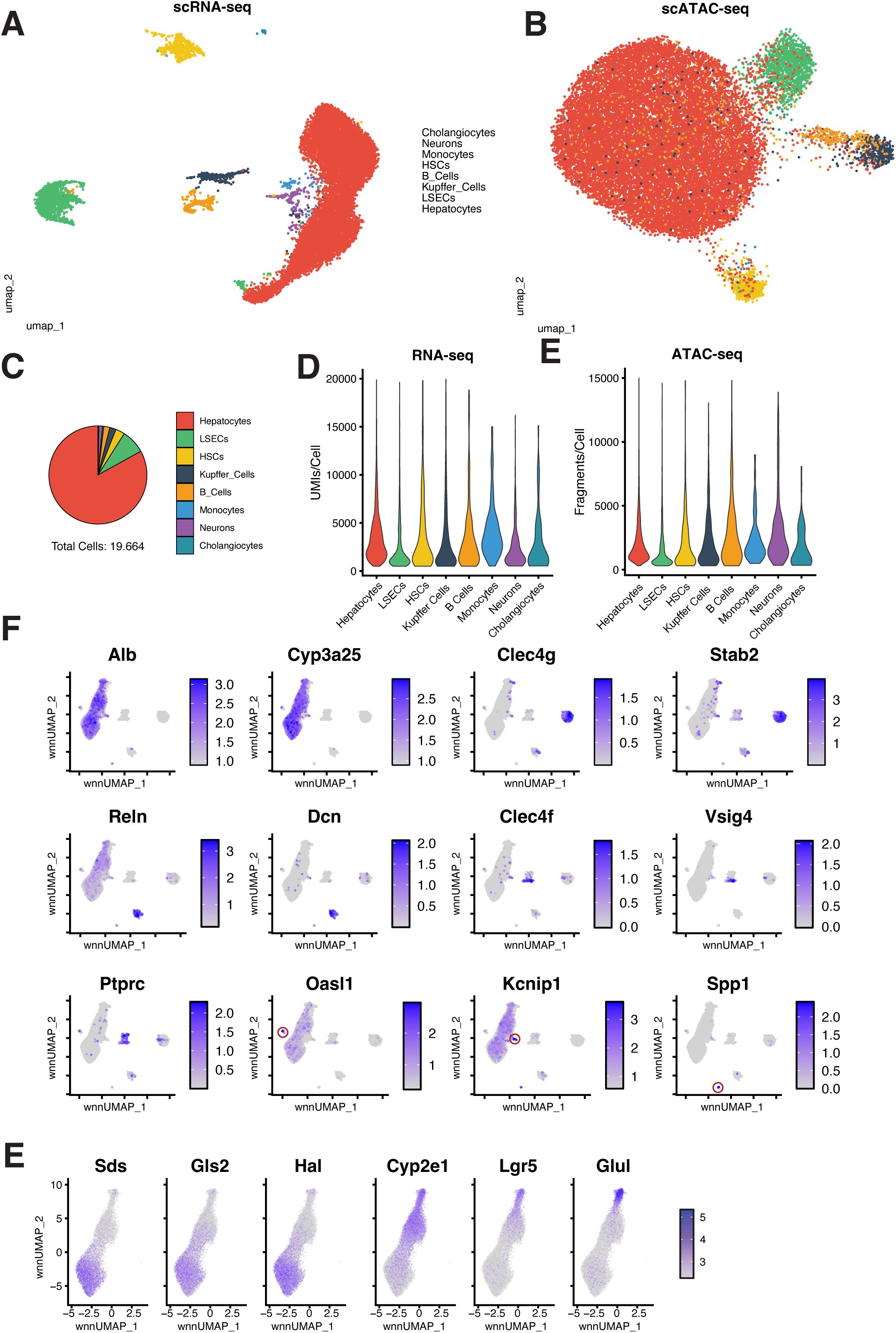
easySHAREseq robustly separates cell types. **(A)** UMAP visualisation of total merged and integrated liver nuclei snRNA-seq data. Nuclei are coloured according to their cell type identified in Fig. 2. (**B**) UMAP visualisation of total merged and integrated liver nuclei snATAC-seq data. Nuclei are coloured according to their cell type identified in Fig. 2. (**C**) Fraction of each recovered cell type relative to all 19,664 nuclei. (**D**) Violin plots depicting the distribution of Unique Molecular Identifiers (UMIs; transcripts) per cell split by cell type. (**E**) Violin plots depicting the distribution of unique fragments per cell split by cell type. (**F**) WNN-UMAPs of 19,664 liver nuclei with nuclei coloured according to the mean expression strength of marker genes for several cell types (*Cyp3a25*: Hepatocytes, *Stab2*: Liver Sinusoidal Endothelial Cells (LSECs), *Reln*: Hepatic Stellate Cells (HSCs), *Clec4f*: Kupffer Cells, *Ptprc*: BCells, *Oasl1*: Monocytes, *Kcnip1*: Neurons *Spp1*: Cholangiocytes). Red circles indicate the position of the cell population showing elevated expression for this marker gene.

**Supplementary Figure 3:**
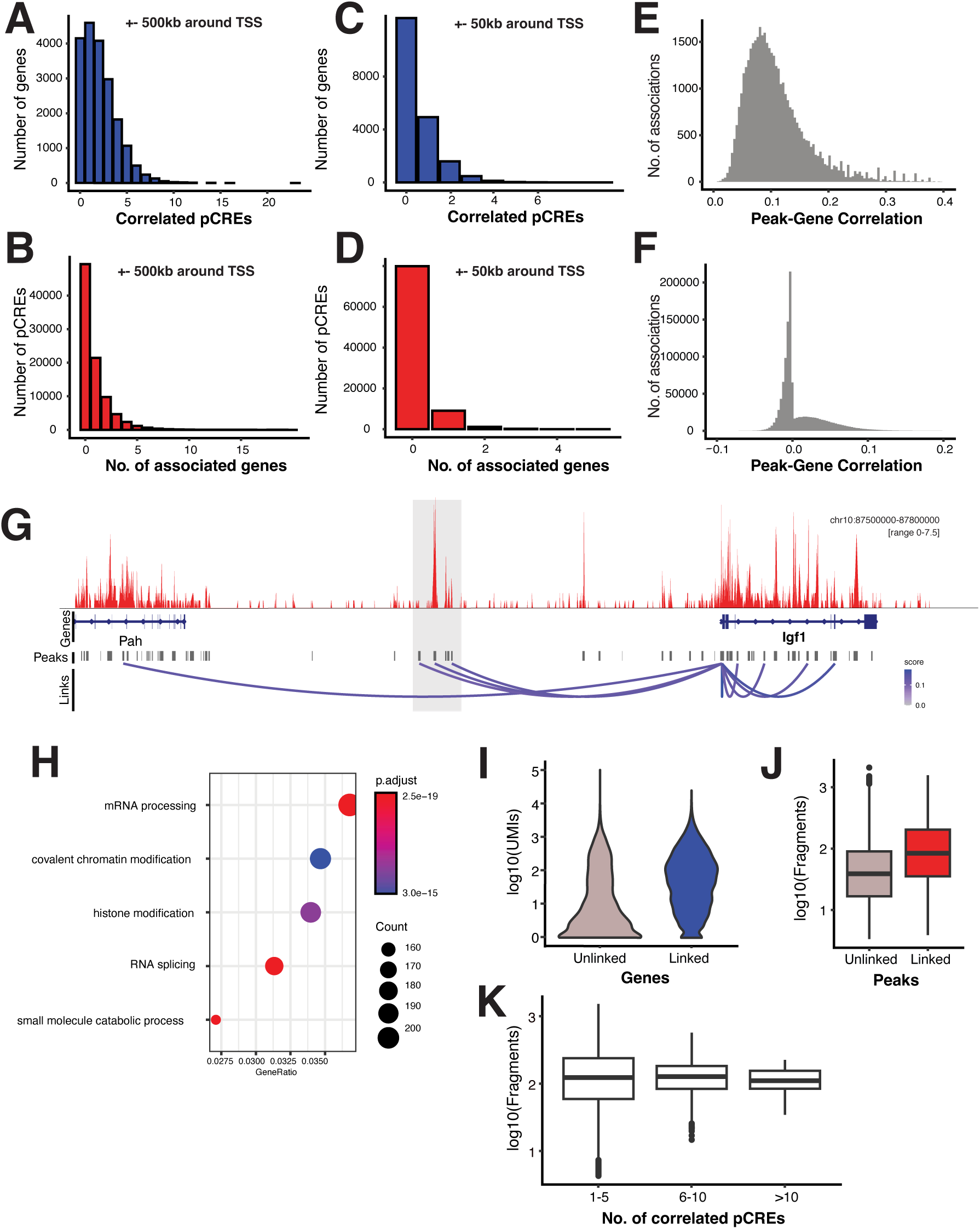
Summary of peak–gene correlations. **(A)** Number of significantly correlated putative cis-regulatory elements (pCREs) per gene (P < 0.05, FDR <= 0.1), considering all peaks ±500kbp of the transcription start site (TSS). (**B**) Number of genes a given pCREs is significantly correlated with (P < 0.05, FDR <= 0.1), considering all peaks ±500kbp of the TSS. (**C**) Number of significantly correlated pCREs per gene (P < 0.05, FDR <= 0.1), considering all peaks ±50kbp of the TSS. (**D**) Number of genes a given pCREs is significantly correlated with (P < 0.05, FDR <= 0.1), considering all peaks ±50kbp of the TSS. (**E**) Histogram of Spearman correlations of all significant peak–gene correlations (P < 0.05). (**F**) Histogram of Spearman correlations of all non-significant peak–gene correlations (P > 0.05). (**G**) Aggregate snATAC-seq track in Liver Sinusoidal Endothelial Cells (LSECs) at the *Igf1* locus and its upstream region. Grey bars indicate open chromatin regions. Loops denote significantly correlated pCREs with *Igf1* and are coloured by their respective Spearman correlation. Shaded grey area denotes a potentially LSEC-specific cis-regulatory element regulating *Igf1* expression. (**H**) Gene Ontology enrichment analysis of genes whose associated pCREs are associated with five or more genes. (**I**) UMIs per gene for genes without and with at least one significant peak-gene correlation (67 vs. 221 mean UMIs/gene; p < 2.2e^-16^; two-sided Welch’s t-test) (**J**) Fragments per peak for peaks with and without a correlated gene (72 vs. 158 mean fragments/gene; p < 2.2e^-16^; two-sided Welch’s t-test) (**K**) Fragments per peak for pCREs associated with genes with 1-5, 6-10 or more than 10 correlated pCREs (Pearson correlation for transcript vs. fragment counts of linked pairs: r=0.004; p = 0.39)

## Methods

### Animal Model & Tissue preparation

#### Mice

All animal experimental procedures were carried out under the licence number EB 01-21M at the Friedrich Miescher Laboratory of the Max Planck Society in Tübingen, Germany. The procedures were reviewed and approved by the Regierungspräsidium Tübingen, Germany. Liver was collected from both male and female wild-type C57BL/6 and PWD/PhJ mice aged between 9 to 11 weeks.

#### Study design

From each strain, we generated easySHARE-seq libraries for one male and one female mice (four total). For each individual, we sequenced two sub-libraries, resulting in 8 easySHARE-seq libraries.

#### Cell Culture

For the species-mixing experiment, HEK Cells were cultured in media containing DMEM/F-12 with GlutaMAX™ Supplement, 10% FBS and 1% Penicillin-Streptomycin (PenStrep) at 37°C and 5% CO2. Cells were harvested on the day of the experiment by simply pipetting them off the plate and were then spun down for 5 min at 250G.

Murine OP9-DL4 cells were cultured in alpha-MEM medium containing 5% FBS and 1% PenStrep. On the day of the experiment, the cells were harvested by aspirating the media and adding 4 ml of Trypsin, followed by an incubation at 37°C for 5 min. Then, 5ml of media was added and cells were spun down for 5 min at 250G. After counting both cell lines using TrypanBlue and the Evos Countess II, equal cell numbers were mixed.

#### Liver Nuclei

The liver was extracted, rinsed in HBSS, cut into small pieces, frozen in liquid nitrogen and stored in the freezer at −80 °C for a maximum of two weeks. On the day of the experiment, 1 ml of ice cold Lysis Solution (0.1% Triton-X 100, 1mM DTT, 10mM Tris-HCl pH8, 0.1mM EDTA, 3mM Mg(Ac)2, 3mM CaCl2 and 0.32M sucrose) was added to the tube. The cell suspension was transferred to a pre-cooled Douncer and dounced 10x using Pestle A (loose) and 15x using Pestle B (tight). The solution was added to a thick wall ultracentrifuge tube on ice and topped up with 4ml ice cold Lysis Solution. Then 9 ml of Sucrose solution (10mM Tris-HCl pH8.0, 3mM Mg(Ac)2, 3mM DTT, 1.8M sucrose) was carefully pipetted to the bottom of the tube to create a sucrose cushion. Samples were spun in a pre-cooled ultracentrifuge with a SW-28 rotor at 24,400rpm for 1.5 hours at 4 °C. Afterwards, all supernatant was carefully aspirated so as not to dislodge the pellet at the bottom and 1 ml ice cold DEPC-treated water supplemented with 10µl SUPERase & 15µl Recombinant RNase Inhibitor was added. Without resuspending, the tube was kept on ice for 20 min. The pellet was then resuspended by pipetting ~15 times slowly up and down followed by a 40 µm cell straining step. Counting of the nuclei using DAPI and the Evos Countess II was immediately followed up by fixation.

### easySHARE-seq protocol

#### Preparing the barcoding oligonucleotides

There are two barcoding rounds in easySHARE-seq with 192 unique barcodes distributed across two 96-well plates in each round (see **Suppl. Table 1** for a full list of oligonucleotide sequences). Each barcode (BC) is pre-annealed as a DNA duplex for improved stability. The first round of barcodes contains two single-stranded linker sequences at its ends as well as a 5’ phosphate group to ligate the different barcodes together. The first single-stranded overhang links the barcode to a complementary overhang at the 5’ end of the cDNA molecule or transposed DNA molecule, which originates either from the RT primer or the Tn5 adapter. The second overhang (3bp) is used to ligate it to the second round of barcodes (**Fig.1B**). Each duplex needs to be annealed prior to cellular barcoding, preferably on the day of the experiment. No blocking oligos are needed.

The Round1 BC plates contain 10µl of 4µM duplexes in each well and Round2 BC plates contain 10µl of 6µM barcode duplexes in each well, all in Annealing Buffer (10mM Tris pH8.0, 1mM EDTA, 30mM KCl). Pre-aliquoted barcoding plates can be stored at −20 °C for at least three months. On the day of the experiment, the oligo plates were thawed and annealed by heating plates to 95 °C for 2 min, followed by cooling down the plates to 20 °C at a rate of −2 °C per minute. Finally, the plates were spun down. Until the annealed barcoding plates are needed, they should be kept on ice or in the fridge.

This barcoding scheme is very flexible and currently supports a throughput of ~350,000 cells (assuming 96 indexing primers) per experiment, limited only by sequencing cost and availability of indexing primer. The barcodes were designed to have at least a Hamming distance of 2. See *Supplementary Notes* for further details on the barcoding system and flexibility.

#### Tn5 preparation

Tn5 was expressed in-house as previously described^77^. Two differently loaded Tn5 are needed for easySHARE-seq, one for the tagmentation, loaded with an adapter for attaching the first barcodes (termed Tn5-B2S), and one for library preparation with a standard illumina sequencing adapter (termed Tn5-A-only). See Supplementary Table 1 for all sequences.

To assemble Tn5-B2S, two DNA duplexes were annealed: 20 µM Tn5-A oligo with 22 µM Tn5-reverse and 20 µM Tn5-B2S with 22 µM Tn5-reverse, all in 50 mM NaCl and 10mM Tris pH8.0. Oligos were annealed by heating the solution to 95 °C for 30 s and cooling it down to 20 °C at a rate of 2 °C/min. An equal volume of duplexes was pooled and then 200 µl of unassembled Tn5 was mixed with 16.5 µl of duplex mix. The Tn5 was then incubated at 37 °C for 1 hour, followed by 4 °C overnight. The Tn5 can then be stored at −20 °C. In our hands, Tn5 did not show a decrease in activity after 10 months of storage.

To assemble Tn5-A-only, 10 µM of Tn5-A and 10.5 µM Tn5-reverse was annealed using the same conditions as described above. Again, 200 µl of unassembled Tn5 was mixed with 16.5 µl of Tn5-A duplex and incubated at 37 °C for 1 hour, followed by 4 °C overnight. The Tn5 can then be stored for later and repeated use for more than 10 months at −20 °C. We observed an increase in all Tn5 activity during the first months of storage, possibly due to continued transposome assembly in storage.

#### Fixation

One million liver nuclei (“cells” for short) were added to ice-cold PBS for 4 ml total. After mixing, 87 µl 16% formaldehyde solution (0.35%; for liver nuclei) or 25 µl 16% formaldehyde solution (0.1%; for HEK and OP9 cells) was added and the suspension was mixed by pipetting up and down exactly 3 times with a P1000 pipette set to 700 µl. The suspension was incubated at room temperature for 10 min. Fixation was stopped by adding ice-cold Stop-Mix (224 µl 2.5M glycine, 200 µl 1M Tris-HCl pH8.0, 53 µl 7.5% BSA in PBS). The suspension was mixed exactly 3 times with a P1000 pipette set to 850 µl and incubated on ice for 3 min followed by a centrifugation at 500G for 5 min at 4°C. Supernatant was removed and the pellet was resuspended in 1 ml Nuclei Isolation Buffer (NIB; 10mM Tris pH8.0, 10mM NaCl, 2mM MgCl2, 0.1% NP-40) and kept on ice for 3 min followed by straining the suspension with a 40 µm cell strainer. It was then spun down at 500G for 3 min at 4°C and re-suspended in ~100-200µl PBSi (1x PBS + 0.4 U/µl Recombinant RNaseInhibitor, 0.04% BSA, 0.2 U/µl SUPERase, freshly added), depending on the amount of input cells. Cells were then counted using DAPI and the Countess II and concentration was adjusted to 2M cells/ml using PBSi.

#### Tagmentation

In a typical easySHARE-seq experiment for this study, 8 tagmentation reactions with 10,000 cells each followed by 3 total RT reactions were performed. This results in sequencing libraries for around 30,000 cells. To increase throughput, simply increase the amount of tagmentation and RT reactions accordingly. No adjustment is needed to the barcoding. Each tube and PCR strip until the step of Reverse Crosslinking was coated before use by rinsing it with PBS+0.5% BSA to maximize cell recovery. Even though all cells could theoretically be tagmented or reverse transcribed in one big reaction, we found separating them into smaller volumes resulted in better data quality.

For each tagmentation reaction, 5 µl of 5X TAPS-Buffer, 0.25µl 10% Tween, 0.25µl 1% Digitonin, 3 µl PBS, 1 µl Recombinant RNaseInhibitor and 9µl of H2O was mixed. TAPS Buffer was made by first making a 1M TAPS stock solution in H2O, followed by adjustment of the pH to 8.5 by titrating 10M NaOH. Then, 4.25ml H2O, 500µl 1M TAPS pH8.5, 250µl 1M MgCl2 and 5ml N-N-Di-Methyl-Formamide (DMF) was mixed on ice and in order. When adding DMF, the buffer heats up so it is important to be kept on ice. The resulting 5X TAPS-Buffer can then be stored at 4°C for short term use (1-2 months) or for long-term storage at −20°C (> 6 months). Then, 5 µl of cell suspension at 2M cells/ml in PBSi was added to the tagmentation mix for each reaction, mixed thoroughly and finally 1.5µl of Tn5-B2S was added. The reaction was incubated on a shaker at 37°C for 30 min at 850 rpm. Afterwards, all reactions were pooled on ice into a pre-cooled 15ml tube. The reaction wells were washed with ~30 µl PBSi which was then added to the pooled suspension in order to maximize cell recovery. The suspension was then spun down at 500G for 3 min at 4°C. Supernatant was aspirated and the cells were washed with 200µl NIB followed by another centrifugation at 500G for 3 min at 4°C.

We only observed cell pellets when centrifuging after fixation and only when using cell lines as input material. Therefore, when aspirating supernatant at any step it is preferable to leave around 20-30µl liquid in the tube. Additionally, it is recommended to pipette gently at any step as to not damage and fracture the cells.

#### Reverse Transcription

As stated above, three tagmentation reactions were combined into one RT reaction. When increasing cells to more than 30,000 per RT reaction, we observed a steep drop in reaction efficiency. The Master Mix for one RT reaction contained 3µl 100µM RT-primer, 2µl 10mM dNTPs, 6µl 5X MaximaH RT Buffer, 4.5µl 50% PEG6000, 1.5 µl H2O, 1.5µl SUPERase and 1.66µl MaximaH RT. The RT primer contains a polyT tail, a 10bp UMI sequence, a biotin molecule and an adapter sequence used for ligating onto the first round of barcoding oligos. The cell suspension was resuspended in 10µl NIB per RT reaction and added to the Master Mix for a total of 30µl. As PEG is present, it is necessary to pipette ~30 times up and down to ensure proper mixing. The RT reaction was performed in a PCR cycler with the following protocol: 52°C for 12 min; then 2 cycles of 8°C for 12s, 15°C for 45s, 20°C for 45s, 30°C for 30s, 42°C for 2min and 50°C for 3 min. Finally, the reaction was incubated at 52°C for 5 more minutes. All reactions were then pooled on ice into a pre-cooled and coated 15ml tube and the reaction wells were washed with ~40µl NIB, which was then added to the pooled cell suspension in order to maximise cell recovery. The suspension was then spun down at 500G for 3 min at 4°C. Supernatant was aspirated and the cells were washed in 150µl NIB and spun down again at 500G for 3min at 4°C. This washing step was repeated once more, followed by resuspension of the cells in 2ml Ligation Mix (400µl 10x T4-Buffer, 40µl 10% Tween-20, 1460µl Annealing Buffer and 100µl T4 DNA Ligase, added last).

#### Single-cell barcoding

Using a P20 pipette, 10µl of cell suspension in the ligation mix was added to each well of the two annealed Round1 BC plates, taking care as to not touch the liquid at the bottom of each well. The plates were then sealed, shaken gently by hand and quickly spun down (~ 8s) followed by an incubation on a shaker at 25°C for 30 min at 350 rpm. After 30 min, the cells from each well were pooled into a coated PCR strip using a P200 multichannel pipette set to 30µl. In order to pool, each row was pipetted up and down three times before adding the liquid to a PCR strip. After 8 columns were pooled into the strip, the suspension was transferred into a coated 5ml tube on ice. This process was repeated until both plates were pooled, taking care to aspirate most liquid from the plates. The cell suspension was then spun down for 3min at 500G at 4°C. Supernatant was aspirated and the cells were resuspended thoroughly in 2 ml new Ligation Mix. Now, 10µl of cell suspension was added into each well of the annealed Round2 barcoding plates using a P20 pipette, taking care as to not touch the liquid within each well. The plates were sealed, shaken gently by hand and spun down quickly followed by incubating them on a shaker at 25°C for 45 min at 350 rpm. The cells were then pooled again using the above-described procedure into a new coated 15ml tube. The cells were spun down at 500G for 3 min at 4°C. Supernatant was aspirated, the cells were washed with 150µl NIB and spun down again. Finally, the cells were resuspended in ~60µl NIB (depending on total amount of cells) and counted. For counting, 5µl of cells were mixed with 5µl of NIB and 1x DAPI and counted on the Evos Countess II, taking the dilution into account. Sub-libraries of 3,500 cells were made and the volume was adjusted to 25µl by addition of NIB.

Using 3,500 cells results in a doublet rate of ~6.3%. The recovery rate of cells after sequencing depends on the input material (and QC thresholds), with cell lines recovering around 80% of input cells (~2,800-3,000 cells) and liver nuclei around 70% (~2,300-2,500 cells).

#### Reverse-Crosslinking

To each sub-library of 3,500 cells, 30µl 2x Reverse Crosslinking (RC) Buffer (0.4% SDS, 100mM NaCl, 100mM Tris pH8.0) as well as 5µl ProteinaseK was added. The sub-libraries were mixed and incubated on a shaker at 62°C for one hour at 800 rpm. Afterwards, they were transferred to a PCR cycler into a deep well module set to 62°C (lid to 80°C) for an additional hour. Afterwards, each sub-library was incubated at 80°C for 10 min and finally 5µl of 10% Tween-20 to quench the SDS and 35µl of NIB was added for a total volume of 100µl. The lysates can be stored at this point at −20°C for at least two days, which greatly simplifies handling many sub-libraries at once. Longer storage has not been extensively tested.

#### Streptavidin Pull-Down

Each transcript contains a biotin molecule as the RT primers are biotinylated, which is used to separate the scATAC-seq libraries from the scRNA-seq libraries. For each sublibrary, 50µl M280 Streptavidin beads were washed three times with 100µl B&W Buffer (5mM Tris pH8.0, 1M NaCl, 0.5mM EDTA) supplemented with 0.05% Tween-20, using a magnetic stand. Afterwards, the beads were resuspended in 100µl 2x B&W Buffer and added to the sublibrary, which were then shaken at 25°C for one hour at 900 rpm. Now all cDNA molecules are attached to the beads whereas transposed molecules are within the supernatant. The lysate was put on a magnetic stand to separate supernatant and beads. It likely is possible to reduce the number of M280 beads in this step, significantly reducing overall costs. However, this has not been extensively tested.

#### scATAC-seq library preparation

The supernatant from each sub-library was cleaned up with a Qiagen MinElute Kit and eluted twice into 30µl 10mM Tris pH8.0 total. PCR Mix containing 10µl 5X Q5 Reaction Buffer, 1µl 10mM dNTPs, 2µl 10µM i7-TruSeq-long primer, 2µl 10µM Nextera N5XX Indexing primer, 4.5µl H2O and 0.5µl Q5 Polymerase was added (All Oligo sequences in Suppl. Table 1). Importantly, in order to distinguish the samples, each sub-library needs to be indexed with a different N5XX Indexing primer. The fragments were amplified with the following protocol: 72°C for 6 min, 98°C for 1 min, then cycles of 98°C for 10s, 66°C for 20s and 72°C for 45s followed by a final incubation at 72°C for 2 min. The number of PCR cycles strongly depends on input material (Liver: 17 PCR cycles, Cell Lines: 15 PCR cycles). The reactions were then cleaned up with custom size selection beads with 0.55X as upper cutoff and 1.4X as lower cutoff and eluted into 25µl 10mM Tris pH8.0. Libraries were quantified using the Qubit HS dsDNA Quantification Kit and run on the Agilent 2100 bioanalyzer with a High Sensitivity DNA Kit.

#### cDNA library preparation

The beads containing the cDNA molecules were washed three times with 200µl B&W Buffer supplemented with 0.05% Tween-20 before being resuspended in 100µl 10mM Tris ph8.0 and transferred into a new PCR strip. The strip was put on a magnet and the supernatant was aspirated. The beads were then resuspended in 50µl Template Switch Reaction Mix: 10µl 5X MaximaH RT Buffer, 2µl 100µM TS-oligo, 5µl 10mM dNTPs, 3µl Enzymatics RNaseIn, 15µl 50% PEG6000, 14µl H2O and 1.25µl MaximaH RT. The sample was mixed well and incubated at 25°C for 30 min followed by an incubation at 42°C for 90 min. The beads were then washed with 100µl 10mM Tris while the strip was on a magnet and resuspended in 60µl H2O. To each well, 40µl PCR Mix was added containing 20µl 5X Q5 Reaction Buffer, 4µl 10µM i7-Tru-Seq-long primer, 4µl 10µM Nextera N5XX Indexing primer, 2µl 10mM dNTPs, 9µl H2O and 2µl Q5 Polymerase. The resulting mix can be split into two 50µl PCR reactions or run in one 100µl reaction. The PCR involved initial incubation at 98°C for 1 min followed by PCR cycles of 98°C for 10s, 66°C for 20s and 72°C for 3 min with a final incubation at 72°C for 5 min. Importantly, in order to distinguish the samples, each sub-library needs to be indexed with a different N5XX Indexing primer. The number of PCR cycles strongly depends on input material (Liver: 15 cycles, Cell lines: 13 cycles).

The PCR reactions were cleaned up with custom size selection beads using 0.7X as a lower cutoff (70µl) and eluted into 25µl 10mM Tris pH8.0. The cDNA libraries were quantified using the Qubit HS dsDNA Quantification Kit.

#### scRNA-seq library preparation

As the cDNA molecules are too long for sequencing (mean length > 700bp), they need to be shortened on one side. To achieve this, 25ng of each cDNA library was transferred to a new strip and volume was adjusted to 20µl using H2O. Then 5µl 5X TAPS Buffer and 0.8µl Tn5-A-only was added and the sample was incubated at 55°C for 10 min. To stop the reaction, 25µl 1% SDS was added followed by another incubation at 55°C for 10 min. The sample was then cleaned up with custom size selection beads using a ratio of 1.3X and eluted into 30µl. Then 20µl PCR mix was added containing 10µl 5X Q5 reaction buffer, 1µl 10mM dNTPs, 2µl 10µM i7-Tru-Seq-long primer, 2µl 10µM Nextera N5XX Indexing primer (note: each sample needs to receive the same index primer as was used in the cDNA library preparation), 4.5µl H2O and 0.5µl Q5 Polymerase. The PCR reaction was carried out with the following protocol: 72°C for 6 min, 98°C for 1 min, followed by 5 cycles of 98°C for 10s, 66°C for 20s and 72°C for 45s with a final incubation at 72°C for 2 min. Libraries were purified using custom size selection beads with a ratio of 0.5X as an upper cutoff and 0.8X as a lower cutoff. The final scRNA-seq libraries were quantified using the Qubit HS dsDNA Quantification Kit and run on the Agilent 2100 bioanalyzer with a High Sensitivity DNA Kit.

#### Sequencing

Both scATAC-seq and scRNA-seq libraries were sequenced simultaneously as they were indexed with different Index2 indices (N5XX). All libraries were sequenced on the Nova-seq 6000 platform (Illumina) using S4 2×150bp v1.5 kits (Read 1: 150 cycles, Index 1: 17 cycles, Index 2: 8 cycles, Read 2: 150 cycles). Libraries were partially multiplexed with standard Illumina sequencing libraries.

#### Custom Size selection beads

To make custom size selection beads, we washed 1ml of SpeedBeads on a magnetic stand in 1ml of 10mM Tris-HCl pH8.0 and re-suspended them in 50ml Bead Buffer (9g PEG8000, 7.3g NaCl, 500ul 1M Tris HCl pH8.0, 100ul 0.5M EDTA, add water to 50ml). The beads don’t differ in their functionality from other commercially available ready-to-use size selection beads. They can be stored at 4°C for > 3 months.

### Analysis

#### Gene annotations and Genomic variants

The reference genome and the Ensembl gene annotation of the C57BL/6J genome (mm10) were downloaded from Ensembl (Version GRCm38, release 102). Gene annotations for PWD/PhJ mice were downloaded from Ensembl. A consensus gene annotation set in mm10 coordinates was constructed by filtering for genes present in both gene annotations.

#### easySHARE-RNA-seq pre-processing

Fastq files were demultiplexed using a custom C-script, allowing one mismatch within each barcode segment. The reads were trimmed using cutadapt^78^. UMIs were then extracted from bases 1-10 in Read 2 using UMI-Tools^79^ and added to the read name. Only reads with TTTTT at the bases 11-15 of Read 2 were kept (> 96%), allowing one mismatch. Lastly, the barcode was also moved to the read name.

#### Species-Mixing Experiments

RNA-seq reads were aligned to a composite hg38-mm10 genome using STAR^80^. The resulting bamfile was then filtered for uniquely mapping reads and reads mapping to chrM, chrY or unmapped scaffolds or containing unplaced barcodes were removed. Finally, the reads were deduplicated using UMItools^79^. ATAC-seq reads were also aligned to a composite genome using bwa^81^. Duplicates were removed with Picard tools and reads mapping to chrM, chrY or unmapped scaffolds were filtered out. Additionally, reads that were improperly paired or had an alignment quality < 30 were also removed. The reads were then split depending on which genome they mapped to and reads per barcode were counted. Barcodes needed to be associated with at least 700 fragments and 500 UMIs in order to be considered a cell for the analysis. A barcode was considered a doublet when either the proportion of UMIs or fragments assigned to a species was less than 75%. This cutoff was chosen to mitigate possible mapping bias within the data.

#### easySHARE-RNA-seq processing and read alignment

We only used Read 1 for all our RNA-seq analyses as we did not need the additional genetic information for this particular analysis. Each sample was mapped to mm10 using the twopass mode in STAR^80^ with the parameters --outFilterMultimapNmax 20 --outFilterMismatchNmax 15. We then processed the bamfiles further by moving the UMI and barcode from the read name to a bam flag, filtering out multimapping reads and reads without a definitive barcode. To determine if a read overlapped a transcript, we used featureCounts from the subread package^82^. UMI-Tools was used to collapse the UMIs of aligned reads, allowing for one mismatch and de-duplication of the reads. Finally, (single-cell) count matrices were created also using UMI-Tools.

#### easySHARE-ATAC-seq pre-processing and read alignment

Fastq files were demultiplexed using a custom C-script, allowing one mismatch within each barcode segment. The paired reads were trimmed using cutadapt^78^ and the resulting reads were mapped to the mm10 genome using bwa mem^81^. Reads with alignment quality < Q30, unmapped, undetermined barcode, or mapped to chrM were discarded. Duplicates were removed using Picard tools. Open chromatin regions were called by subsampling the bamfiles from all samples to a common depth, merging them into a pooled bamfile and using the peak caller MACS2^83^ with the parameters -nomodel -keep-dup -min-length 100. The count matrices as well as the FRiP score was generated using featureCounts from the Subread package^82^.

#### Filtering, Integration & Dimensional reduction of scRNAseq data

The count matrices were loaded into Seurat^84^ and cells were then filtered for >200 detected genes, >500 UMIs and < 20.000 UMIs. The sub-libraries coming from the same experiment were then merged together and normalised. Merged experiments from the same species (one from male mouse, one from female mouse) were then integrated by first using SCTransform^85^ to normalise the data, then finding common features between the two experiments using FindIntegrationAnchors() and finally integrated using IntegrateData(). Lastly, the integrated datasets from C57BL/6 and PWD/PhJ were again integrated using IntegrateData(). To visualise the data, we projected the cells into 2D space by UMAP using the first 30 principal components and identified clusters using FindClusters().

#### Filtering, Integration & Dimensional reduction of scATACseq data

Fragments per cell were counted using sinto and the resulting fragment file was loaded into Signac^86^ alongside the count matrices and the peakset. We calculated basic QC statistics using base Signac and cells were then filtered for a FRiP score of at least 0.3, > 300 fragments, < 15.000 fragments, a TSS enrichment > 2 and a nucleosome signal < 4. Again, sub-libraries coming from the same experiment were merged. We then integrated all four experiments (C57BL/6 & PWD/PhJ, one male & one female mouse each) by finding common features across datasets using FindIntegrationAnchors() using PCs 2:30 and then integrating the data using IntegrateEmbeddings(). To visualise the data, we projected the cells into 2D space by UMAP.

#### Weighted-Nearest-Neighbor (WNN) Analysis & Cell type identification

In order to use data from both modalities simultaneously, we created a multimodal Seurat object and used WNN^42^ clustering to visualise and leverage both modalities for downstream analysis. Afterwards, we assigned cell cycle scores and excluded clusters consisting of nuclei solely in the G2M-phase (2 clusters, 121 nuclei total). Cell types were assigned via expression of previously known marker genes, which allows subsetting the data into cell types. To estimate ambient RNA contamination, we used decontX^87^ supplying the raw count matrix and cell type identities. For subsequent analysis of cell-type specific gene expression and UMAP-plots, we used the de-contaminated counts provided by decontX.

#### Calculating Peak–Gene Associations

Peak–gene associations were calculated following the framework described by Ma et al^28^. In short, Spearman correlation was calculated for every peak–gene pair within a +-500kb window around the TSS of the expressed gene. To obtain a background estimation, we used chromVAR^88^ (getBackgroundPeaks()) to generate 100 background peaks matched in GC bias and chromatin accessibility but randomly distributed throughout the genome. We calculated the Spearman correlation between every background-gene comparison, resulting in a null distribution with known population mean and standard deviation. We then calculated the z-score for the peak–gene pair in question ((correlation − population mean)/ standard deviation) and used a one-sided z-test to determine the p-value. This functionality is also implemented in Signac under the function LinkPeaks(). Increasing the number of background peaks to 200, 350 or 500 for each peak–gene pair does not impact the results (data not shown).

#### Analysis of LSEC zonation markers

To analyse gene expression and chromatin accessibility along LSEC zonation, we ordered LSECs along pseudotime in Monocle3^89^. The seurat object for LSECs was converted into a Monocle cell data set (as.cell_data_set()), clustered as a UMAP (cluster_cells()) followed by graph learning (learn_graph()). Cells with the highest *Wnt2* expression were set as root for pseudotime ordering (order_cells()). Gene expression and Chromatin Accessibility for marker genes was smoothed over pseudotime with local polynomial regression fitting (loess). To identify novel marker genes, we excluded genes with low expression, divided the pseudotime into 10 bins and calculated the moving average (for three bins) across them. We then required the moving average to continuously decrease (for pericentral marker genes) or increase (for periportal marker genes), allowing two exceptions. Lastly, we divided the means for each gene by their maximum to normalise the values. Identification of cis-regulatory elements displaying zonation effects had equal requirements.

#### Gene Ontology Analysis

Gene Ontology Analysis was done using the R package clusterProfiler^90^ with standard parameters.

#### Comparison to external datasets

In order to compare the performance of easySHARE-seq to other multiomic and single-channel technologies, we downloaded the raw data from Martin et al.^36^ (sciRNA-seq3; GEO: GSE186824, first four samples, 9.974 murine E16.5 embryonic cells), Chen et al.^33^ (SNARE-seq, GEO: GSE126074, first 4 replicates, 2.621 adult mouse brain cortex cells), Cao et al.^30^ (sciCAR, GEO:GSE117089, 5968 murine kidney cells), Cusanovich et al.^38^ (sciATAC-seq, GEO: GSE111586, 7.023 murine liver nuclei), Gonzales-Blas et al.^34^ (10x Multiome ATAC+RNA; GEO: GSE218468, data generated with 10x protocol; 7.518 murine liver nuclei) and Nikopolou et al.^37^ (10x scATAC-seq, E-MTAB: E-MTAB-12706, 7.810 murine liver nuclei). We downsampled all datasets to a common sequencing depth (22.000 reads/cell for the RNA-seq, 34.000 reads/cell for the ATAC-seq) and processed them equally. To determine fragments in peaks per cell, we used the peaksets provided by the authors. Su et al.^35^ (10x 3’ expression, 82.168 murine liver cells) did not provide all raw data needed for downsampling and we therefore used the authors provided count matrix. They report a sequencing depth of ~ 180.000 reads / cell. Gonzales-Blas (10x Multiome,7.518 murine liver nuclei) did not report all files necessary for downsampling the ATAC-seq dataset. We therefore used the count matrix as provided. Unfortunately, Ma et al.^28^ (SHARE-seq, 42.948 murine skin cells) did neither report full raw data, code nor a sequencing depth. We therefore used the authors count matrix as provided.

## Data Availability

The easySHARE-seq data reported in this paper can be downloaded with the accession number GSE256434. All code used in data analysis is available at https://github.com/vosoltys/easySHARE_seq.git.

## Acknowledgements

We thank members of the Chan and Jones lab for helpful discussions and critical reading of the manuscript. We are very grateful to Arnar Breevoort and Alex Pollen for sharing tissue preparation protocols and a very helpful research visit. We thank Sinja Mattes and all animal care takers at the Friedrich Miescher Laboratory for their work. We also thank the Genome Center in the Max Planck Institute for Biology Tübingen for providing support. The OP9-DL4 cells were a kind gift from Juan Carlos Zúñiga-Pflücker. M.P. is supported by an International Max Planck Research School fellowship. M.K. and Y.F.C. were supported by the European Research Council Starting Grant 639096 “HybridMiX” and Proof-of-Concept Grant 101069216 “Haplotagging”. The research was supported by the Max Planck Society.

## Author Contributions

V.S. and Y.F.C. designed the experiments. V.S. and M.P. developed the barcoding framework for easySHARE-seq. V.S. developed the rest of the protocol and performed experiments. V.S. performed the computational analyses advised by Y.F.C. V.S. drafted the manuscript. M.P., D.S., M.K. and Y.F.C. helped with experimental or computational support. All authors reviewed the manuscript.

## Declaration of Interest

The authors declare no competing interests.

## Notes

https://www.ncbi.nlm.nih.gov/geo/query/acc.cgi?acc=GSE256434

### Competing Interest Statement

The authors have declared no competing interest.

### Summary of Updates

Slight revision of the discussion section and Supplementary Notes.

