## Supplementary Notes for "Flexible and high-throughput simultaneous profiling of gene expression and chromatin accessibility in single cells"

### Flexibility and Applicability of the easySHARE-seq framework

EasySHARE-seq uses a flexible barcoding framework that can be tailored to various experimental designs. As mentioned in the main text, it allows for sequencing of fragment lengths of > 200bp, which can be critical in e.g. studies investigating patterns of allele-specific expression or profiling of individual cancer cells and their mutations. However, in study designs not dependent on SNP Coverage, sequencing costs can be cut with no downside by only sequencing 100bp per fragment.

The entire barcoding can also be easily adapted into other protocols, such as scTCR-seq (CITR-seq; unpublished), allowing for paired investigation of T-cell receptor chains in millions of cells.

It is also straightforward to adapt easySHARE-seq to a scRNA-seq only protocol with equal or even increased throughput as well as sample indexing, allowing to run a single experiment for e.g. multiple replicates. To achieve this, the tagmentation step can simply be skipped and the RT-primer can be switched out for /5Phos/GGGCTCGGAGATGTGTATAAGAGACAGNNNNNNNNNN-[8bp-Sample-Index]-/biodT/TTTTTTTTTTTTTTTTTTTTTTTTTVN. Before the first 10bp UMI in R2 a 8bp barcode was introduced which can represent an individual sample, timepoint, replicate or simply cell pool. A separate reverse transcription reaction(s) for each sample with a differently barcoded RT-primer for each of them needs to be performed. Afterwards, all cells can be pooled and the protocol can be performed as described. Additionally, the amount of cells per sub-library and thus throughput can be increased linearly depending on how many different sample barcodes (RT-primers) are used. For example, using 2 RT-primers with different sample barcodes, the amount of cells per sublibrary can be doubled. PCR cycles need to be adjusted for the increased cell numbers. Altogether, this allows to upscale easySHARE-seq several fold relatively straightforward with minor additional reagent costs or processing time. Throughput of easySHARE-seq is only limited by the availability of Nextera N5XX Indexing Primers, theoretically enabling the simultaneous profiling of up to a million cells.

Switching to a scATAC-seq only protocol is done by simply excluding the RT step as well as the Streptavidin pull-down. This will significantly cut experimental cost as no RT, RNase Inhibitors or Streptavidin beads are required, bringing the cost per cell down to ~2.5 cents/cell (in a 100.000 cell experiment). Furthermore, fixation strengths can be decreased which in our hands led to improved data quality.

However, upscaling the scATAC-seq part is less straightforward since in contrast to the RNA-seq, any additional barcodes cannot be within the insert and must be in the index reads. Currently, the barcoding scheme is designed to span the least number of bases possible which involves ‘breaking up’ the Read2 sequencing primer site between the barcoding oligos and the Tn5/RT-primer. Introducing barcoded Tn5’s with additional barcodes would thus entail either (1) forgoing sample indexing on Index2 and thus be not suitable for upscaling this part or (2) significantly increase the number of bases sequenced in Index1 and thus decrease one of the key strengths of our protocol. Additionally, this is complicated by the need to change the sequence of the barcoding oligos as well as the need for unassembled Tn5.

Lastly, to cut further costs on easySHARE-seq, it is possible to perform only a single ligation step. Leaving out the first ligation (in the BC plates 1) still produces easySHARE-seq libraries as the initial overhang is 8bp long and therefore can theoretically form a stable hybridization at room temperature.

### Critical Optimization steps to use easySHARE-seq efficiently

The general molecular steps of easySHARE-seq are quite robust. However, in order to use easySHARE-seq efficiently, some prior optimizations should be performed.
As with most scRNA-seq experiments, sample preparation and fixation have the highest impact on success and quality of the experiment. As sample preparation can be quite different between tissues, general good practice is including a sufficient amount of RNase Inhibitor, especially when input material is concentrated in small volumes.
The strength of fixation has a direct impact on data quality of both the scATAC-seq and scRNA-seq. In general, higher fixation leads to an increase of data quality in the scRNA-seq but makes the tagmentation in the scATAC-seq less efficient. Therefore, fixation parameters can to some extent be adjusted based on the requirements and importance of the respective output modality. Fixation strength should also be optimized in a tissue-specific manner. For example, fixing cell lines in 0.15% PFA was generally sufficient for data quality and maintaining cell integrity throughout the protocol. Liver nuclei needed a higher fixation of 0.35% and Bone Marrow Cells (*not shown*) needed to be fixed in 1% PFA as they are both fragile and contain low amounts of mRNA molecules. Another critical factor is the fixation volume, e.g. fixation with 1% PFA in 1 ml leads to a different outcome than fixation in 4 ml. Generally, it is advisable to fix input material in higher volumes and with a low concentration of cells (~1M/ml) as this leads to more consistent results and less clumping. For initial experiments to devise fixation strength, we advise to simply skip the barcoding step. This can be done by using a standard Tn5 for ATAC-seq and the following RT-primer: GTCTCGTGGGCTCGGAGATGTGTATAAGAGACAG/biodT/TTTTTTTTTTTTTTTTTTTTTTTTTVN.
Using these modifications ensures that both the scATAC-seq and scRNA-seq can be amplified with standard Nextera N7XX and N5XX primers, allowing for cost-efficient testing of easySHARE-seq parameters. Additionally, cell integrity should be periodically checked to detect cell clumps and assess cell integrity.
Another critical aspect is minimizing freeze-thaw cycles for barcoding oligos, especially for oligos containing phosphorylation modifications. Repeated freezing and thawing leads to a strong decline in protocol efficiency. Please feel free to contact the First Author for further questions.

### Example workflow of easySHARE-seq library generation of 200.000 cells

To perform an experiment with a yield of ~200.000 cells, one needs to perform ~48 tagmentation reactions with 10.000 cells per reaction. After tagmentation, those get distributed into 16 RT reactions. Barcoding is then performed as described in one reaction. Afterwards, 96 sublibraries of ~3.500 cells are aliquoted and can be further processed. After Reverse Crosslinking, the samples can be stored at -20C until the next day.
To simplify the cleanup of the lysate for the scATAC-seq library preparation, they can be cleaned-up with size selection beads by adding 150ul per well.

### Discussion of ambient RNA contamination results

As seen in **Suppl. Fig. 1O**, decontX identifies mean contaminated counts of 9.6% and median contaminated counts of 1.4%. The authors of decontX report mean contamination values of 1-4% in commercial droplet-based protocols and 11-14% in plate-based protocols

[1], suggesting that easySHARE-seq performs better than other plate-based assays but does not reach the results of protocols that physically separate the cells. These values also suggest that few cells that are heavily contaminated strongly increase the overall estimation of contaminated counts. This could be explained by doublets and/or wrongly assigned cell types since cell type information is passed onto decontX and builds the foundation for identifying contamination.
Overall, we found that our analyses are robust to decontamination. We thus see this filtering step as not strictly necessary but do recommend it. An alternative approach to using decontX employed in other studies is filtering out UMIs that are only associated in one sequencing read [2].

### Potential alterations to the protocol in order to improve ATAC-seq data quality

Compared to the original protocol, ATAC-seq quality is decreased, potentially resulting in less resolution and variance in downstream analyses. We see multiple potential aspects of the protocol that when changed might improve ATAC-seq data quality.
The greatest potential for improvement likely lies in the fixation step as lower fixation strength generally yields higher data quality. Therefore, a simple comparison of data quality in e.g., 0.1% vs. 0.2% PFA-fixated cells of the same cell type would be informative. That said, exact fixation strength is highly cell- and tissue-type dependent.
A second aspect to explore is the tagmentation step. Since fixation reduces Tn5 cutting efficiency, increasing tagmentation temperature or duration could partially compensate for this. For example, increasing tagmentation reactions to 42°C or 45min could result in improved ATAC-seq data quality, although care must be taken not to compromise cell integrity.

1. Yang, S. *et al.* Decontamination of ambient RNA in single-cell RNA-seq with DecontX. *Genome Biol* **21**, 57 (2020).
2. Ma, S. *et al.* Chromatin Potential Identified by Shared Single-Cell Profiling of RNA and Chromatin. *Cell* **183**, 1103-1116.e20 (2020).
